# Remodelling of chromatin architecture and super-enhancer landscape in lamin A/C depleted HGSOC

**DOI:** 10.64898/2026.09.07.749802

**Authors:** Shreyasi Dey Sarkar, Poulami Goswami, Amrutayani Panda, Vishnu S. Mishra, Bhabani S. Mohanty, Roli Budhwar, Kakoli Bose, Shuvojit Paul, Ayan Banerjee, Pradip Chaudhari, Kaushik Sengupta

## Abstract

Lamins are nuclear intermediate filament proteins that maintain nuclear architecture through interactions with the chromatin. The A- and B-type lamins tether the genome at the peripheral lamina underlying the inner nuclear membrane in the form of heterochromatic lamina-associated domains (LADs). However, LADs associated with A-type lamins are confined not only to lamina but also to the nuclear core, thereby pointing to their distinct and multifarious roles in chromatin organisation and regulation. Previous studies involving lamin B1 depletion depicted the detachment of LADs from the nuclear periphery, accompanied by alteration of chromatin distribution while preserving the topologically associating domains (TADs) in structurally intact form. In this piece of work, we have shown, for the first time in HGSOC, the effects of lamin A/C knockdown on chromatin organisation, which was characterised by significant detethering of gene-poor chromatin from the periphery along with active (A) to inactive (B) compartment switching. This was associated with a rewiring of oncogenic super-enhancer elements corroborated by the differential gene expression profile. Overall analysis tipped the scale in favour of reduced cellular proliferation upon lamin A/C knockdown. This finding was strengthened by the observed proliferative potential of spheroids ex vivo and tumours in a mouse xenograft model.

**Graphical Abstract:** Figure-
Schematic illustration showing the effect of lamin A/C knockdown in HGSOC on higher-order chromatin organisation and transcriptional dynamics, leading to reduced proliferation

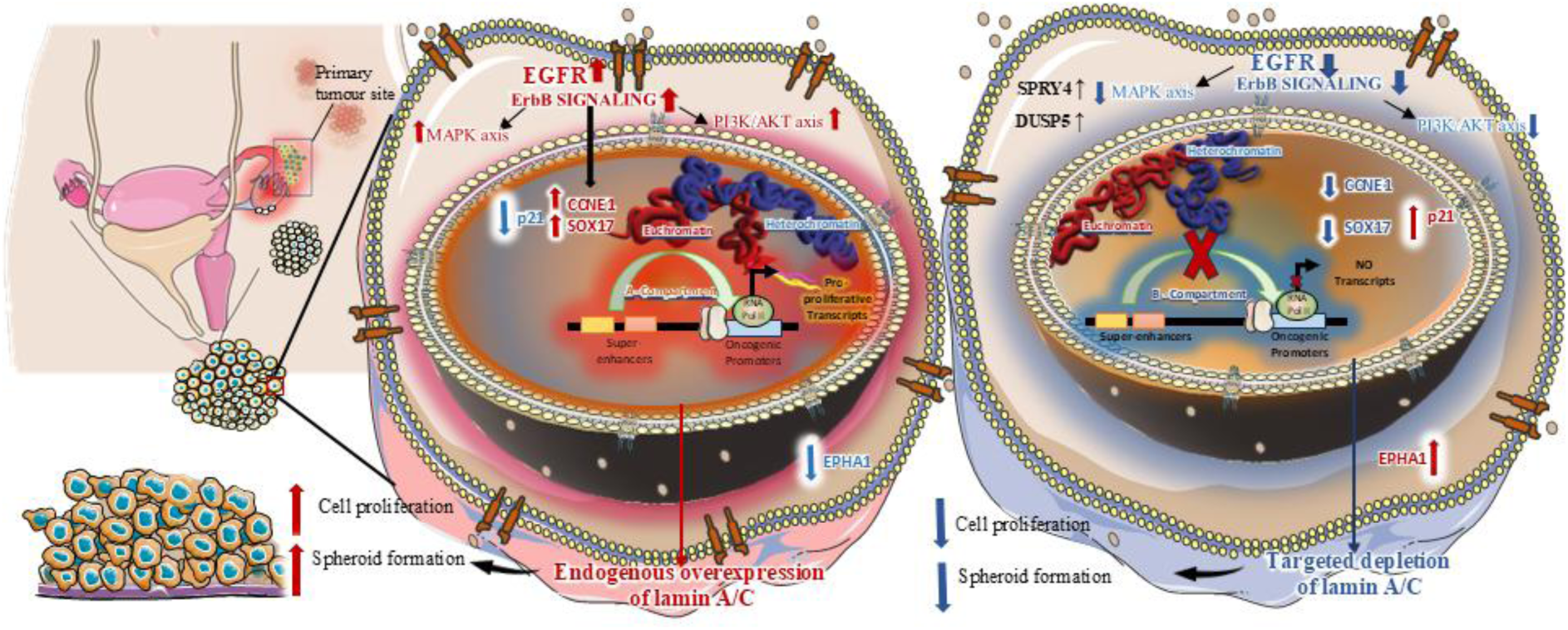

## Introduction

Ovarian cancer is one of the most lethal gynaecological malignancies, ranking eighth in female cancer incidence and second in gynaecological cancer mortality worldwide (1). High-grade serous ovarian cancer (HGSOC) represents the most common and lethal histotype of epithelial ovarian cancer. Largely due to late-stage diagnosis, HGSOC shows dismal long-term survival outcomes, with 5-year relative survival rates remaining below 30–35% (2).

Amongst different drivers modulating the malignancy, expression of lamins, the type V intermediate filament proteins, has been reported widely. The four lamin genes in vertebrates, LMNA, LMNB1, LMNB2, and LIII, encode four major and three minor types of lamin proteins, the major ones being B1, B2, A, and C. Amongst these, A-type lamins include splice variants-lamin A and lamin C, collectively called lamin A/C and encoded by the LMNA gene. A-type lamins not only contribute to forming the nuclear lamina but also form a nucleoplasmic scaffold, unlike B-type lamins, which are essentially peripheral. Besides providing mechanical integrity to the nucleus, both A- and B-type lamins contribute to establishing the 3D chromosomal landscape by formation of lamina-associated domains (LADs) (3), which are enriched in repressive heterochromatic histone marks and represent transcriptionally silent states at the nuclear periphery. The peripheral scaffold of lamin A/C can directly interact with core histones and DNA via the C-terminal tail and Ig-fold domain, facilitating direct chromatin docking at the periphery. It interacts with Emerin, LAP2alpha, beta, and MAN1 (the LEM Domain Proteins) (4), which recruit Barrier-to-Autointegration Factor (BAF) (5) to bridge DNA to the lamina meshwork. A recent study demonstrates that lamin A/C interacts with PRR14 and tethers it to the nuclear periphery, where it recruits HP1 to repressed heterochromatin regions (6). In the nuclear interior, lamin A/C does not form a dense meshwork; rather maintains a mobile, low-assembly state by associating with LAP2alpha (Lamina-Associated Polypeptide 2alpha), which escorts it to open, actively transcribed chromatin (7–9). The complete genomic regions physically bound by nuclear lamins were termed as lamin-Interacting Domains (LiDs) (10). Sonication-derived LMNA-LiDs are comprised of compacted, gene-poor heterochromatin domains at the nuclear periphery that can resist nuclease digestion. MNase-derived LMNA-LiDs capture nuclease-accessible, gene-dense regions enriched in active/facultative marks (with open euchromatin features), often reflecting interactions driven by the soluble nucleoplasmic pool of lamin A/C complexed with LAP2alpha. Nucleoplasmic lamin A/C localises into dynamic internal foci and speckles that interact with transcriptionally active compartments and spatial splicing domains to modulate RNA synthesis and overall transcription, regulating cellular proliferation (11, 12). LADs consist of one or more whole TADs (Topologically Associating Domains), and physical repositioning of chromatin frequently occurs in discrete TAD-sized units. Inter-TAD higher-order contacts and radial repositioning of TAD units are contributed by the lamin A/C pool (13). The TADs segregate into distinct active (A) or inactive (B) chromatin compartments, in 3D physical space relative to the nuclear periphery, depending on specific DNA methylation profiles and additional epigenetic marks. This leads to differential recruitment of transcription factors in these A and B compartments with distinct transcriptional outcomes, indirectly modulated by lamin expression (14). The role of lamin A/C in cancer is context-dependent, and its effects on tumorigenesis are largely debated to date(15–17). Compared to normal tissue, lamin A/C expression is distinctly dysregulated among tumour types (15, 16, 18), and this expression profile is often regarded as a mechanical trade-off between migratory invasion and cell survival in the context of malignant transformation (19).

In several cancers, like gastric carcinoma (20), pancreatic intraductal papillary neoplasms (21), lung cancer (22–24) pediatric tumors such as neuroblastoma, osteosarcoma, and Ewing sarcoma, reduced lamin A/C expression has been shown to lead to with low patient survival rate which can be alleviated by lamin A/C re-expression (25–28) whereas lamin A/C is overexpressed in various aggressive cancers such as Colorectal cancer, High-Grade Prostate Cancer and Glioblastoma Multiforme (29) (30), (31). Interestingly, investigations in ovarian cancer have also provided contradictory results with respect to increases and decreases in lamin A expression compared to normal tissue(32–36).

Although cohort-dependent variability and contextual downregulation of lamin A/C have been reported in ovarian cancer (35), high-density protein microarray and immunoblot analysis established lamin A/C to be significantly overexpressed in malignant ovarian tissues relative to non-cancer controls (34).

In NIH-OVCAR3, an aggressive HGSOC model, elevated endogenous lamin A expression not only helps to provide regular mechanobiological support (37, 38) but has also been found to confer essential survival and proliferative advantage by facilitating homologous recombination (HR) and non-homologous end joining (NHEJ) DNA double-strand break repair pathways (39, 40). A significant positive correlation was reported between elevated lamin A/C expression and poorer overall survival in advanced ovarian cancer patients, also unveiling the role of endogenous overexpression of A-type lamins in reduced responsiveness of primary tumour tissues to chemotherapeutic agents (41, 42). Nuclear deformities coupled with an altered epigenetic landscape were found to be associated with dysregulated expression of lamin A/C in ovarian carcinoma (43, 44).

Beyond maintaining structural and mechanical integrity, the lamina functions as a critical platform coordinating the higher-order three-dimensional (3D) architecture of the genome (45, 46). The mobile nucleoplasmic pool of lamin A/C, associating with both hetero- and euchromatic regions, dynamically tunes chromatin accessibility to transcription machinery and Polycomb-mediated repression, also affecting replication timing (9). 3D chromatin positioning, which has been widely reported to be altered in the context of cancer, is significantly regulated by lamin A/C (47–49). Non-random relative positioning of chromosomes in nuclear space has often been found to facilitate the genomic rearrangements between them to initiate malignancy (50). Besides pervasive copy-number alterations and high chromosomal instability (CIN), extensive literature documented dysregulated organisation of higher-order chromatin in ovarian cancer, more so in HGSOC (51–54). Besides large-scale physical alterations in chromosomes, which correlated with poor clinical outcome (55, 56), mounting evidence documented territorial redistribution (55) and compartment switching of the ovarian cancer genome (57–60). The disputed expression and occupancy of CTCF support dysregulation at the level of TADs in ovarian cancer (61, 62). The abrogated insulating boundaries and rearrangement of chromatin loops physically shuffle to favour ectopic contacts between distal regulatory elements and oncogenic promoters, driving aberrant transcriptional activation supporting rapid proliferation (63, 64). During malignant transformation in HGSOC, the master-regulatory transcription factors coalesce at de novo super-enhancers that loop directly to oncogenic drivers and cell-cycle genes. Recent evidence underscores that rewired super-enhancer landscapes and associated transcription factor circuitries directly promote chemoresistant phenotypes in HGSOC (65, 66).

Although the downstream oncogenic consequences and transcription factor networks associated with rewired super-enhancers in ovarian cancer are well documented and established (67, 68), the upstream mechanistic drivers such as nuclear architectural factors and structural scaffolds governing this extensive spatial reorganisation remain largely unexplored. Historically, altered expression of lamin A/C in cancer was primarily viewed in the light of nuclear morphological defects, cytokinesis failure, and numerical chromosomal instability leading to aneuploidy (35, 69); but the active role of overexpressed lamin A/C as a topological coordinator of the 3D cancer genome remains to be charted. Our study bridges this critical gap by demonstrating that elevated lamin A/C acts as a structural scaffold that stabilises oncogenic genome topology. Targeted depletion of A-type lamins triggers extensive spatial rearrangement of peripheral heterochromatin and overall chromatin reorganisation, including prominent compartment switching, which modulates oncogenic super-enhancers and targets the ERBB pathway, along with other tumour-promoting molecular regulators, substantially attenuating HGSOC cell proliferation. Thus, our work connects nuclear envelope biology with 3D epigenomics, establishing lamin A/C as a promising paradigm in understanding the higher-order genomic topology leading to aberrant transcriptional circuitry sustaining aggressive ovarian malignancies.

## Material and methods

### Cell culture

NIH-OVCAR3 cells (hence referred to as WT-OVCAR3) were maintained in ATCC-formulated RPMI-1640 Medium (R6504) supplemented with 1% Penicillin-Streptomycin (Gibco™,15-140-148) and 20% foetal bovine serum (FBS, F4135). Cells were maintained in culture flasks or dishes at 37°C in a humidified incubator containing 5% CO₂. Trypsin-EDTA (0.25%) (Gibco™, 25200056) was used to detach the cells from the flask surface. Cell counting was performed using trypan blue exclusion assay by mixing the cell suspension and trypan blue solution (Gibco™, 15250061) in a 1:1 ratio. Viable cells were counted using an automated cell counter.

### Lentiviral shRNA-mediated knockdown

Stable LMNA-knockdown OVCAR3 cell line (hence referred to as shOVCAR3 cells) was generated using a lentiviral shRNA expression system- lamin A/C shRNA (h) Lentiviral Particles (sc-35776-V), containing expression constructs encoding 19-25 nt (plus hairpin) shRNA designed to knockdown lamin A/C expression. After transduction, stable cells expressing the shRNA were isolated via selection with puromycin (Sigma, P8833). For selection and propagation of successfully transduced cells, a puromycin concentration of 7.5ug/ml was maintained in the culture medium according to the IC50 determined in the laboratory a priori.

### Spontaneous spheroid formation

Budding spheroids were allowed to form spontaneously by culturing both cell lines in their regular growth media, with or without puromycin, on adherent plates. Spheroids formed spontaneously from the cellular monolayer upon reaching maximum confluency as described previously(70).

### Population doubling time assay

WT-OVCAR3 and shOVCAR3 cells were seeded in triplicate into 3 identical 24-well plates to calculate population doubling time (PDT) at kinetic time points of 24 hours, 72 hours, and 120 hours. Cells were seeded in triplicate wells at an initial density of 3x10^4^ cells per well. Each of the three plates was harvested at distinct intervals of 24 hours, 72 hours, and 120 hours post-seeding. While assessing well viability at each indicated time point, the designated plate was removed from incubation and processed for cell counting. Adherent cells were detached by regular trypsinisation. Cell suspensions were gently pipetted to ensure complete dissociation of cell aggregates. Viable cells were quantified via Trypan Blue dye exclusion using an automated cell counter. Total viable cell numbers per well were recorded in triplicate. Calculation of Population Doubling Time (PDT) was done in the prime log phase between 24 hours and 72 hours in WT-OVCAR3, as follows:

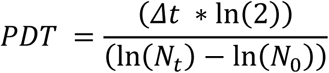

Where *Δt* is the duration of the experiment: 24 and 72 hours; *N_t_* is the cell count at the final time point of 72 hours, and *N*_0_ is the cell count at the initial time point of 24 hours.

### Wound healing assay

A cell-free gap was created using a Culture-Insert 2 Well (Ibidi, 80209), which provides two separate cell-seeding areas. Two such inserts were used to grow WT-OVCAR3 and shOVCAR3 cells under standard culture conditions until a confluent monolayer was formed within the insert chambers. Once the cells reached confluency, the insert was carefully removed, generating a defined cell-free gap between adjacent cell populations on two sides. Immediately after insert removal, phase-contrast images of the gap were captured using an inverted microscope (0 h time point). Thenceforth, the cells were maintained under standard culture conditions to allow regular growth and subsequent closure of the cell-free gap. Images of the gaps, acquired at designated time intervals (0 hours, 12 hours, 24 hours, 36 hours, 48 hours), were used to monitor the gap closure and compare the proliferative capacity of cells. The surface area of gaps was quantified by ImageJ software (ImageJ bundled with 64-bit Java 1.8.0_172). Statistical analyses were performed using GraphPad Prism (v8.4.2). Data are presented as mean±SEM from three independent biological replicates. Differences in percentage wound closure between WT-OVCAR3 and shOVCAR3 across time points (0–48 hours) were evaluated using a two-way ANOVA followed by Šídák’s post-hoc multiple comparisons test. Statistical significance was defined as p < 0.05.

### Quantitative real-time PCR

QIAGEN RNAeasy mini kit (74104, Qiagen) was used to isolate RNA from cell pellets. Purity and Integrity of RNA were analysed by running the isolated RNA samples on FA gel. 5000 ng RNA was used to prepare cDNA using a cDNA synthesis kit (K1622, Thermo Fisher Scientific, Waltham, MA, USA) with oligodT primers according to the manufacturer’s protocol. Quantitative real-time PCR and data analysis were performed in the Applied Biosystems QuantStudio 5 Real-Time PCR System using primer sequences as mentioned earlier (39).

### Western blotting

Protein concentration was estimated by Bradford assay, and western blotting was performed as described earlier (39). Primary antibodies used in this study were Anti- Lamin A/C Monoclonal Antibody HRP (MA5-46598, Thermo Fisher, USA), and Anti-GAPDH antibody (ab9485, Abcam, USA) at 1:1000 and 1:500 dilutions, respectively. The secondary antibody used was HRP-conjugated Goat-Anti-Rabbit (32460, Thermo Scientific, USA). ImageJ software (ImageJ bundled with 64-bit Java 1.8.0_172) was used to perform densitometric analysis of Western blot bands. Signal intensities of lamin A/C were normalised to the corresponding loading control of GAPDH for each lane. Relative fold changes were calculated by normalising the knockdown group to the wild-type. Statistical analyses across independent biological replicates (n = 3) were conducted using GraphPad Prism (v8.4.2) by performing a two-tailed one-sample t-test. Data are presented as mean±SD, and p < 0.05 was considered statistically significant.

### Immunofluorescence staining

Cells were grown on sterile coverslips for immunostaining. On reaching 70-80% confluency, the coverslips were processed and subsequently mounted as described earlier(71). Primary antibody dilution for Histone H3K9me3 antibody (D4W1U, Cell Signalling Technology, USA), H3K27me3 antibody (AB_2636821, Active motif), H3K4me3 antibody (39159, Active motif) and H3K36me3 antibody (61101, Active motif) were 1:500. Primary antibody dilution for Anti- Lamin A/C Monoclonal Antibody HRP (MA5-46598, Thermo Fisher, USA) was 1:100. Secondary antibodies conjugated with Alexa Fluor 546 and Alexa Fluor 488 were used at a dilution of 1:400.

### Confocal imaging and image analysis

For confocal imaging, the slides were visualized by 100X oil DIC N2 objective 1.40 NA/1.515 RI in NIKON TiE inverted microscope with a 4X digital zoom. The images were captured in resonant mode. The excitation filters used were 450/50, 525/50, 595/50, and the first dichroic mirror used was 405/488/561. The lasers used were Multi line Argon-Krypton mixed gas laser λ488nm, solid state laser λ405nm and solid-state laser λ561nm. Images were processed using Ni Elements Analysis AR Ver 4.13 and ImageJ software (ImageJ bundled with 64-bit Java 1.8.0_172). For 3D-FISH imaging, the resonant scanner with a scanner zoom of 6.0 and line averaging of 4.0 was used. While capturing the images, the pinhole was maintained at 69.0 um; a one-way scan direction and channel series mode line 1->4 was used. The Z-stack imaging was carried out in a step size of 0.15- 0.2 µm. Quantitative image analysis calculating mean fluorescent intensity of nuclear proteins was carried out using ImageJ software (ImageJ bundled with 64-bit Java 1.8.0_172). Statistical analysis was performed using GraphPad Prism (v8.4.2). Differences between the WT-OVCAR3 and shOVCAR3 were evaluated using a two-tailed unpaired t-test with Welch’s correction for unequal variances. Data are presented as mean±SD, and P-value < 0.05 was considered statistically significant. Imaris 7.7.2 was used for 3D surface rendering in volumetric confocal stacks of 3D-FISH.

### Sample preparation for RNA sequencing

Cells resuspended in TRIzol samples were stored at -80°C until further processing. All the samples were quantified using Qubit RNA BR Assay (Invitrogen, Cat# Q10211). RNA purity was checked using NanoDrop, and RNA integrity was checked on TapeStation using RNA Screen Tapes (Agilent, Cat# 5067-5576). After confirmation, QC-passing samples were taken for library preparation and sequencing. HyperPrep Kit (Roche KAPA, Cat # 0000141759) protocol was used to prepare libraries for mRNA sequencing. An initial concentration of 200ng of total RNA was taken for the assay. mRNA molecules were captured using magnetic oligo Poly(T) beads (Roche KAPA, Cat#7962240001). Following purification, the enriched mRNA was fragmented using divalent cations at elevated temperature (94°C for 6 minutes). The cleaved RNA fragments were copied into first-strand cDNA using reverse transcriptase enzyme. In a combined reaction, second-strand cDNA synthesis and A-tailing were carried out, which converts the cDNA: RNA hybrid to double-stranded cDNA (dscDNA), incorporates dUTP into the second cDNA strand, and adds dAMP to the 3’ ends of the resulting dscDNA. This was followed by Adapter Ligation, in which dsDNA indexing adapters with 3’ dTMP overhangs were ligated to library insert fragments. The adapter-ligated products were then purified and enriched using high-fidelity, low-bias PCR (The strand marked with dUTP was not amplified, allowing strand-specific sequencing) using the following thermal conditions: initial denaturation at 98°C for 45 sec; 13 cycles of 98°C for 15 sec, 60°C for 30 sec, 72°C for 30 sec; final extension at 72°C for 1 min. PCR products were then purified and checked for fragment size distribution on a Fragment Analyser using the dsDNA Reagent Kit (1-6000bp) (Agilent, Cat# 5191-6576). Prepared libraries were quantified using the Qubit HS Assay (Invitrogen, Cat# Q32854). The obtained libraries were pooled and diluted to the final optimal loading concentration. The pooled libraries were then analysed on the Illumina NovaSeqX Plus instrument to generate 7.5GB data per sample.

### RNA sequencing processing and differential gene expression analysis

Bulk RNA sequencing was performed on high-quality RNA samples extracted from the six samples (comprising three shOVCAR3 and WT-OVCAR3 replicates). Stranded RNA sequencing libraries were prepared, yielding 78.7 million to 104.7 million total reads per sample. The quality of the raw and processed reads was assessed using FastQC (v0.12.1) (72) and summarised using MultiQC (version 1.33) (73). GC content was consistent across libraries (46.24–46.71%), while high Phred scores (38.90–38.99) and over 94.23% Q30 bases confirmed good sequencing quality. Raw RNA sequencing data were quality-filtered and adapter-trimmed using fastp (v1.3.1) (74). High-quality paired-end reads were aligned to the human reference genome (GRCh38/hg38) using the alignment tool STAR (75) (v2.7.11b) in two-pass mode; coordinate-sorted BAM files were generated. Gene-level read counts were subsequently generated from the coordinate-sorted BAM files using featureCounts (Subread v2.1.1) (76) by assigning properly paired, non-chimeric fragments to annotated exons (-p – countReadPairs -B -C -t exon -g gene_id) with the GENCODE (v49) primary assembly basic gene annotation. The resulting count matrix was imported into DESeq2 (v1.42.1) (77) in R version 4.5.3 for differential expression analysis. Differential gene expression between shOVCAR3 and WT-OVCAR3 samples was assessed using the Wald test. P values were corrected for multiple hypothesis testing using the Benjamini–Hochberg procedure, and genes with an adjusted P value < 0.05 and an absolute log₂ fold change ≥ 0.585 (corresponding to a 1.5-fold change in expression) were considered differentially expressed. Ensembl gene identifiers were mapped to HGNC gene symbols, gene names, and Entrez Gene identifiers using the org.Hs.eg.db annotation database (v3.14.0) through the AnnotationDbi package (v1.56.2) in R. The annotated gene lists were used for downstream visualisation and functional enrichment analyses. Gene expression patterns were visualised using principal component analysis (PCA) and volcano plots, and hierarchical clustering heatmaps were generated using variance-stabilised transformed (VST) expression values.

Detailed methods for pathway analysis have been described in the supplementary Section.

### 3D-Fluorescent In-Situ Hybridisation (3D-FISH)

3D FISH was performed on cells grown to a confluency of approximately 70% on glass coverslips as mentioned earlier (71). After post-hybridisation and washing, the coverslips were processed for immunostaining with Anti- Lamin A/C Monoclonal Antibody HRP (MA5- 46598, Thermo Fisher, USA) at a dilution of 1:100 and mounted with Vectashield Vibrance (VECTASHIELD, H-1800-2). Slides were stored at 4°C until imaging.

### Experimental workflow and library preparation for Hi-C

Intact WT-OVCAR3 and shOVCAR3 cells were first crosslinked with 1% ice-cold formaldehyde (Sigma, P6148) in PBS to lock spatial chromatin interactions in place, followed by an incubation period of 10 mins at room temperature. Subsequently, ice-cold glycine was added to a final concentration of 125-150 mM to quench the cross-linking reaction. After quenching for 5 mins at room temperature, the cells were covered with ice-cold PBS and collected with a cell scraper, followed by spinning at 3500 × g for 8 mins at 4C. The cell pellets were stored at -80°C for further use. The crosslinked samples were then digested using a restriction enzyme cocktail provided in the Arima-Hi-C Plus Kit (A510008, Arima Genomics). The resulting sticky ends were filled in with biotinylated nucleotides and proximity-ligated to the adjacent ends. After reversing the crosslinks, the purified genomic DNA was mechanically sheared using Covaris S220 to generate an average fragment size of ∼300-400 bp, and enriched for the biotin-labelled ligation junctions using streptavidin pull-down beads. These enriched Hi-C DNA fragments were converted into sequencing libraries using NEBNext® Ultra™ II DNA Library Prep Kit for Illumina (E7645S, New England BioLabs). Library quality and size distribution were verified via standard quality control assays with the TapeStation 2200 System (Agilent) using D1000 ScreenTape (5067–5587) and D1000 sample buffer (5067–5583). Library concentration was measured using a Qubit 4 Fluorometer (Life Technologies) with the Qubit™ 1X dsDNA HS Assay Kits (Q33231, ThermoFisher Scientific). Paired-end sequencing was done on an Illumina platform using the NovaSeq 6000 platform to an average target depth of 300 million reads per sample.

### Hi-C Data preprocessing and analysis

The raw FASTQ sequencing reads were evaluated using FastQC v0.12.1 to check baseline quality scores and look for adapter contamination. Fastp v1.0.1 (74) with default parameters was used to trim adapters and filter out low-quality bases (Phred score < 20). The clean, paired-end reads were processed through the Hi-C-Pro v3.1.0 pipeline (78). The reads were mapped to the human reference genome (hg38) using Bowtie2 v2.5.4, applying sensitive local and global end-to-end alignment settings to properly capture chimeric ligation junctions. To ensure high-confidence mapping, only those read pairs where both anchors showed a mapping quality (MAPQ) score ≥ 10 were kept. Final libraries were assessed by standard QC and sequenced on an Illumina platform to a depth of approximately 300 million reads per sample. The quality of raw Hi-C sequencing data was initially assessed using FastQC v0.12.1 (79). Low-quality reads and potential adapter contamination were removed using Fastp v1.0.1 (74) with default parameters. Filtered reads were then processed using the Hi-C-Pro v3.1.0 pipeline (78) to generate genome-wide contact matrices. Briefly, Hi-C-Pro alignment was performed against the hg38 reference genome using Bowtie2 v2.5.4 (80) with sensitive local and global alignment settings. The reads with a mapping quality ≥ 10 were retained for the downstream analysis. The Hi-C-Pro was configured using default settings to process DpnII-digested fragments, removing singleton, multi-mapped, and duplicate reads. Contact matrices were generated at 50Kb and 1Mb resolution in upper-triangular format and normalised using iterative correction (ICE) with a maximum of 100 iterations. Low-count bins (bottom 2%) were filtered out before downstream analysis. The topologically associated domains (TADs) were identified using ARMATUS v2.3.0 (81). A/B compartments were identified from balanced 1 Mb Hi-C contact matrices using the eigs-cis function implemented in Cooltools v0.7.1 (82). The first eigenvector (E1) orientation was uniformly adjusted so that positive E1 values represented transcriptionally active A compartments and negative E1 values represented inactive B compartments. Eigenvector phasing was established by correlating E1 values with hg38 GC content and GENCODE v49 transcription start site (TSS) derived gene density across 1-Mb bins. For genome-wide compartment analysis, each 1 Mb genomic bin was categorised according to its compartment status in WT and sh-OVCAR3 cells, as stable (A→A, B→B) or switching (A→B, B→A). Changes in compartment organisation were quantified using the difference in compartment eigenvector values (ΔE1 = E1_shOVCAR3_ − E1_WT-OVCAR3_).

To assess whether compartment remodelling correlates with transcriptional changes, differentially expressed genes (DEGs; adjusted p < 0.05, |log2 fold change| ≥ 0.585) from RNA sequencing data were mapped to their corresponding 1-Mb compartment bins by TSS coordinates and assigned to the respective compartment transition class. Differences in expression distributions across compartment classes were evaluated using Mann–Whitney U tests, and genome-wide association between ΔE1 and transcriptional change was determined using Spearman rank correlation.

3D chromatin structures were reconstructed from the Hi-C contact frequencies with established nuclear lamina interaction maps (GSM1376181, GSM1541019, and GSE109924) using Chrom3D v1.0.2 (83) and visualised. Spatial visualisations were realised in UCSF ChimeraX v1.7.1 (84), where Chromosome 18 (orange) and Chromosome 19 (magenta) were rendered inside a standard, uniform spherical nuclear envelope of radius = 6.0 μm, with 85% transparency. Both WT and sh models were set to identical coordinate perspectives (tilt: x = - 25°, y = 35°) to ensure direct, unbiased structural comparison across conditions. Super-enhancer (SE) loci analysis is described in the supplementary methods section.

### In Vivo Study

The animal studies were performed according to the national guidelines for animal handling. The project was approved by the Institutional Animal Ethics Committee of ACTREC (IAEC- 10/2026). WT-OVCAR3 and shOVCAR3 cells were cultured in RPMI medium supplemented with 20% FBS and 1% antibiotic solution and puromycin to a concentration of 7.5ug/ml in the case of shOVCAR3, under standard cell culture conditions. To evaluate the tumourigenic potential of the two cell lines, WT-OVCAR3 and shOVCAR3 cells were subcutaneously inoculated into 6–7-week-old nude mice to establish xenograft tumours. Each mouse received 3 × 10⁶ cells suspended in serum-free RPMI medium and mixed with Matrigel (Corning, CLS354234) in a 1:1 ratio (v/v; to a final concentration of 50%, immediately before injection. Each experimental group consisted of three nude mice.

Animals were monitored for tumour development and growth. Forty-five days post-inoculation, the mice were euthanised, and the tumours were excised. Tumour tissues were fixed in 10% neutral buffered formalin (NBF) for 24 hours before histopathological analysis. After euthanising the mice, tumour tissues were fixed in 10% NBF for 24 hours prior to histopathological analysis. The samples were then subjected to dehydration through a series of graded ethanol (70–100%) treatments, followed by xylene wash, and paraffin wax embedding. Tissue sections of 5 μm thickness were cut and affixed to clean glass slides. Subsequently, the tissue sections were subjected to deparaffinization, rehydration and haematoxylin and eosin (H&E) staining. The stained tissue sections were then dehydrated, xylene-treated, and mounted with DPX mounting medium. The mounted sections were then assessed for histological changes under a light microscope (Labomed, Lx-300, USA).

### Setup and initial conditions of coarse-grained representation of the lamin-chromatin system

A coarse-grained polymer model was used to examine the function of lamin-mediated interactions in chromatin organisation and compartmentalisation (85–92), where chromatin and lamin proteins were depicted as freely diffusible and interacting beads. A single polymer chain made up of 428 distinct chromatin beads that mimic euchromatin and heterochromatin, which represent variations in compaction and activity, was identified as two distinct chromatin states (85, 93). Lamin A and lamin B were considered to be separate, freely diffusing particles. This distinction incorporated experimentally observed variations in function and interaction specificity of lamins (94–96). 400 beads for each of lamin A and lamin B were considered when modelling the WT scenario, whereas lamin A was reduced to 80 and lamin B increased to 600 beads to represent sh conditions. Simulations were performed in a three-dimensional box of dimensions 10*σ* × 10*σ* × 15*σ*, where *σ* denotes the characteristic bead diameter. Chromatin conformations were initialised by using a self-avoiding random-walk (SAW) algorithm where beads were sequentially placed at a fixed step length subject to a minimum separation constraint from all previously placed beads. Backtracking was applied when no valid placement was found within a fixed number of attempts. Heterochromatin identity was assigned to adjacent blocks of 20 beads, with 30% of blocks being designated as heterochromatin and the remaining as euchromatin, reflecting the domain-scale organisation of chromatin rather than a per-bead random assignment (93, 97, 98). Lamin A and lamin B particles were placed independently, with the majority of lamin B particles (95%) being initialised near the upper wall, with the remaining distributed uniformly throughout the domain. Lamin A particles were also placed uniformly. Following the initial placement, configurations were relaxed through (i) an iterative pairwise overlap-removal procedure displacing any particle pair closer than 1.1*σ*, and (ii) an iterative bond-repair step enforcing that all chromatin bond lengths remained within the FENE extension limit *R*_0_ prior to the start of dynamics. Note that the FENE bonding potential has been elucidated in the supplementary section. Interaction strengths *ε_ij_* were then ramped linearly from 10% to 100% of their target values over the first 5000 simulation steps to avoid instabilities arising from residual steric overlaps.

### Interaction potentials

The non-bonded interactions between particles were modelled using a truncated Lennard-Jones (LJ) potential (98, 99),

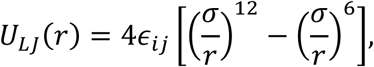

applied for inter-particle distances *r* < *r_c_* and set to zero otherwise. The interaction parameter *ε_ij_* was chosen to encode preferential affinities between different species. The corresponding pairwise force, obtained as *F_ij_* = −*ΔU_LJ_*, is

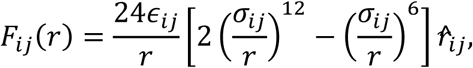

where *r̂_ij_* is the unit vector along the interparticle separation and *σ_ij_* = (*σ_i_* + *σ_j_*)⁄2. Force contributions were truncated at *r* = *r_c_*_,*ij*_ and capped at a maximum magnitude *F_max_* for numerical stability.

The interaction matrix and bonding potentials, along with the methods for confining boundary and nuclear envelope potential, have been detailed in the supplementary section.

### Molecular dynamics simulations and implementation

The system was evolved using Langevin dynamics to account for thermal fluctuations and viscous damping,

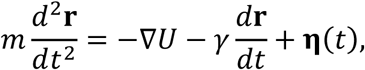

where *γ* is the friction coefficient and **η**(*t*) is a Gaussian random force satisfying the fluctuation-dissipation relation. In practice, the equations of motion were integrated using a velocity-Verlet position update combined with an exact Ornstein-Uhlenbeck (OU) thermostat step for the velocity update (a BAOAB-type Langevin splitting scheme), applied per timestep *Δt* as follows:

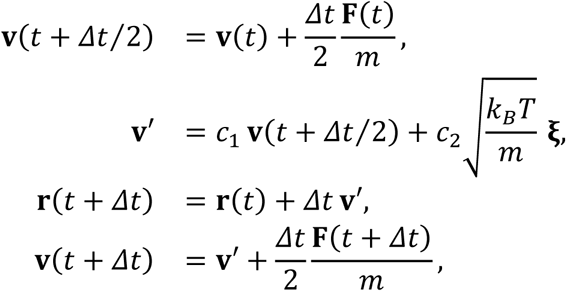

where **ξ** is a vector of independent standard normal random variables, and the OU coefficients are given exactly by

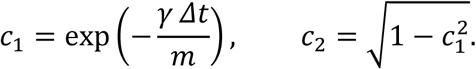

This exact treatment of the friction and stochastic forcing terms avoided the discretisation error associated with an explicit Euler–Maruyama update of the noise term (100). All simulations were implemented in custom MATLAB scripts. The computational workflow included (i) initialisation of particle coordinates, (ii) evaluation of bonded and non-bonded forces, (iii) stochastic integration of equations of motion, and (iv) trajectory storage. Force calculations were optimised using matrix-based operations to ensure computational efficiency. The simulation parameters are elucidated in Table 3.

**Table 3.** Summary of parameters used in coarse-grained simulations.

| Parameter | Symbol | Value |
| --- | --- | --- |
| Chromatin bead diameter | $\sigma$ | 1 |
| Lamin A bead diameter | $\sigma_L$ | $0.5\sigma$ |
| Lamin B bead diameter | $\sigma_{LB}$ | $0.4\sigma$ |
| Box dimensions | $L_x, L_y, L_z$ | $10\sigma \times 10\sigma \times 15\sigma$ |
| Chromatin beads | $N_c$ | 428 |
| Lamin A (WT/KD) | $N_{LA}$ | 400 / 80 |
| Lamin B (WT/KD) | $N_{LB}$ | 400 / 600 |
| Mass (EC, HC, L, LB) | $m$ | 1.0, 1.5, 0.25, 0.13 |
| Friction coeff. (EC, HC, L, LB) | $\gamma$ | 1.0, 1.5, 0.25, 0.13 |
| Global LJ cutoff | $r_{\text{cut,LJ}}$ | $1.8\sigma$ |
| Time step | $\Delta t$ | $1 \times 10^{-4} \tau$ |
| Simulation length | – | $4 \times 10^5$ steps |
| Equilibration ramp | – | 5000 steps (10%→100% $\epsilon$) |

## Results

### Lamin A/C knockdown in OVCAR3

The wild-type OVCAR3 cell line (WT-OVCAR3) was shown to express lamin proteins to a significant extent compared to normal ovarian surface epithelial cells (IOSE) (36), and the elevated level of endogenous lamin A was ascribed to promote HR and NHEJ-mediated DNA repair mechanisms, thereby imparting chemoresistance (39). Taking cues from our previous studies, we aimed to generate a stable lamin A/C knockdown OVCAR3 cell line (shOVCAR3) using a lentiviral shRNA expression system to study the global effect of lamin A/C in facilitating tumourigenesis. We preferred knockdown to knockout to make this study physiologically relevant. Knockdown of lamin A/C was confirmed by Western blot, immunofluorescence and real-time PCR (Fig.1). Quantitative real-time PCR showed a 2.8-fold reduction in the transcript level (Fig 1A), while western blot analysis recorded a 70% reduction at the level of protein expression (Fig. 1B, C). This was further supported by analyses of fluorescence intensity of lamin A/C staining from confocal micrographs (Fig. 1D, E).

**Fig. 1.**
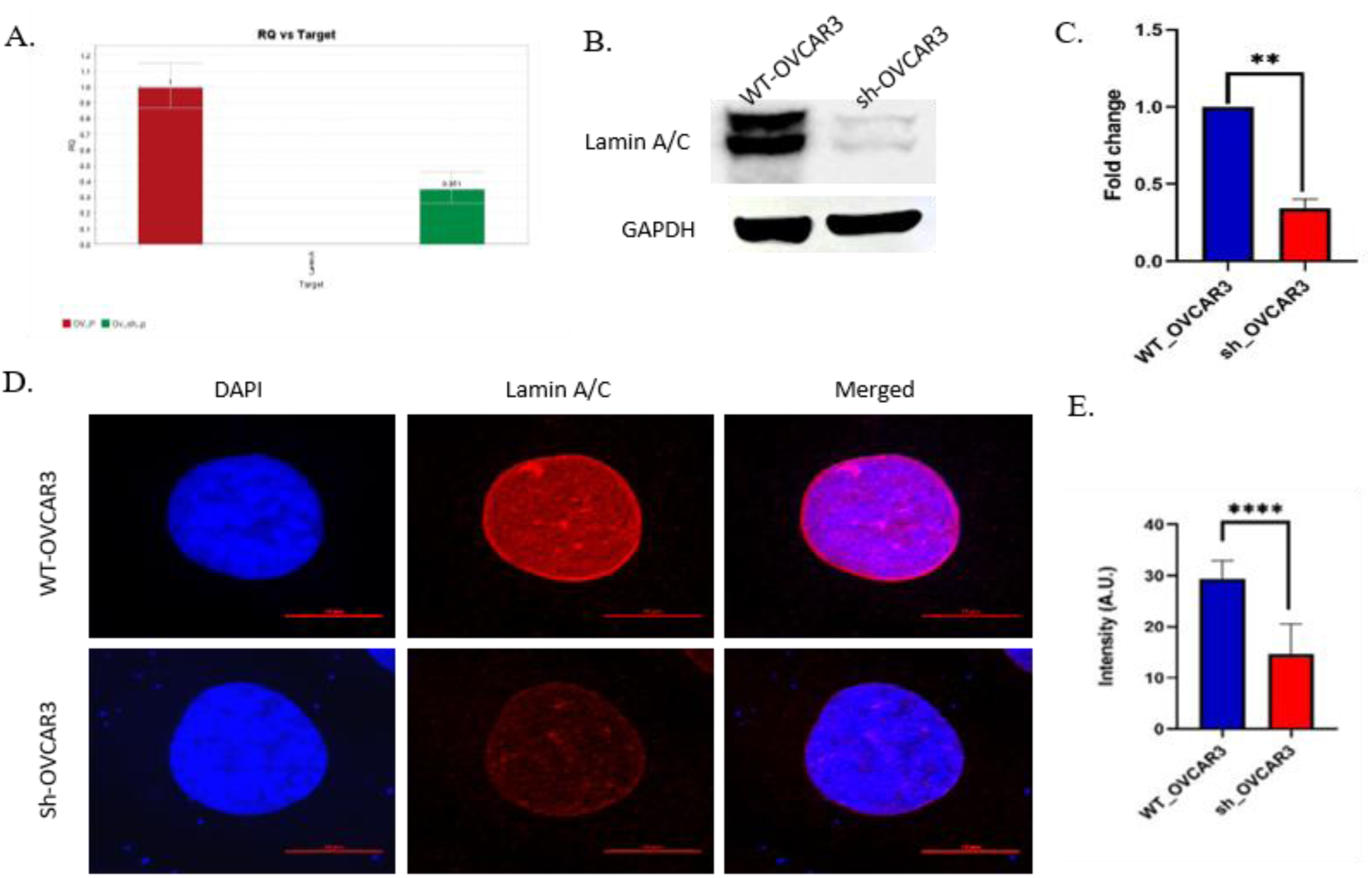
Knockdown of Lamin A/C in OVCAR3 cells. **(A)** mRNA expression of LMNA gene was checked in shOVCAR3 cells in comparison with WT-OVCAR3. **(B)** Western blot showing knockdown of lamin A/C in OVCAR3 cells. **(C)** The expression of lamin A/C in WT and shOVCAR3 cells, quantified from immunoblot done against lamin A/C, normalized to GAPDH. **(D)** Confocal images depicting expression of lamin A/C in WT-OVCAR3 and shOVCAR3 cells (scale bar: 10 μm). **(E)** The expression of lamin A/C in WT and shOVCAR3 cells, quantified from confocal images depicting mean fluorescent intensity (Mean Gray Value) of nuclei.

### Knockdown of lamin A/C causes spatial reorganisation of chromatin in 3D nuclear space

Compared to the normal counterparts, malignant cells were reported to show widely altered chromosomal positioning along with genome instability and genetic abnormalities (101–103). Such remodelled positioning of chromosome territories was also found to cluster oncogenic and metastasis-related genes (104). Chromosome territorial redistribution has remained a major hallmark distinguishing between non-malignant and malignant cells in specific cancers (105, 106). Differential expression of lamins has been widely discussed to alter chromatin architecture in various diseases, including cancer (107). Direct interactions between lamin A/C and chromatin have been elucidated at the docking sites of heterochromatic regions, which are widely referred to as lamina-associated domains (LADs). In addition to its peripheral localisation, the mobile nucleoplasmic pool of lamin A/C also interacts with euchromatic DNA (9, 108). Hence, we asked the most pertinent question here – how does the chromosome landscape change as a sequel to lamin A/C knockdown? To understand this, we chose to invest in studying the 3D orientation of chromosome 18 (Chr. 18) and chromosome 19 (Chr. 19) as representatives of heterochromatin and euchromatin, respectively. We performed 3D fluorescence in situ hybridisation (3D FISH) to examine the nuclear distribution of these two chromosomes. Chromosome 18, residing in the outer nuclear periphery, is a gene-poor, late-replicating chromosome containing repressive domains enriched in repressive heterochromatin marks (109–111). Chromosome 19, on the other hand, having the highest gene density in the human genome, contains euchromatin-rich, transcriptionally active regions and occupies an interior nuclear distribution (110, 112, 113).

In WT-OVCAR3 cells characterised by high endogenous levels of lamin A/C, we noticed a conventional polar-opposite 3D radial distribution within the nucleus, with chromosome 18 being distinctly peripheral (Fig. 2, panel A) and chromosome 19 showing a more central positioning (Fig. 2, panel C). On the contrary, Chr 18 & 19 underwent large-scale spatial redistribution in shOVCAR3. The distinct differential positioning of both chromosomes in WT and shOVCAR3 is evident from the representative 3D-volume snapshots and 3D surface rendition images of the wild-type and knockdown cells depicted in Fig. A. IV- D. IV and A. V- D. V, respectively. Upon lamin A/C depletion, in shOVCAR3 cells we found a partial significant detethering of chromosome 18 from nuclear periphery towards nuclear centre (Fig. 2B). Along with this dislodgement we found a simultaneous but slight change in radial distribution of chromosome 19 from centre towards periphery (Fig. 2D). We quantified the effect of lamin A/C knockdown on repositioning of chromosomes by calculating the radial distribution of the chromosome territories as depicted in the schematic of our model (Fig. 2E). The radial distribution plot (Fig. 2F) clearly elucidated the heterochromatin detethering from periphery to centre concomitantly with peripheral euchromatin redistribution upon loss of lamin A/C in an endogenous overexpression background in HGSOC. We project this to be the first report depicting global genome repositioning of whole chromosomes as a function of lamin A/C expression profile in an HGSOC model.

**Fig. 2.**
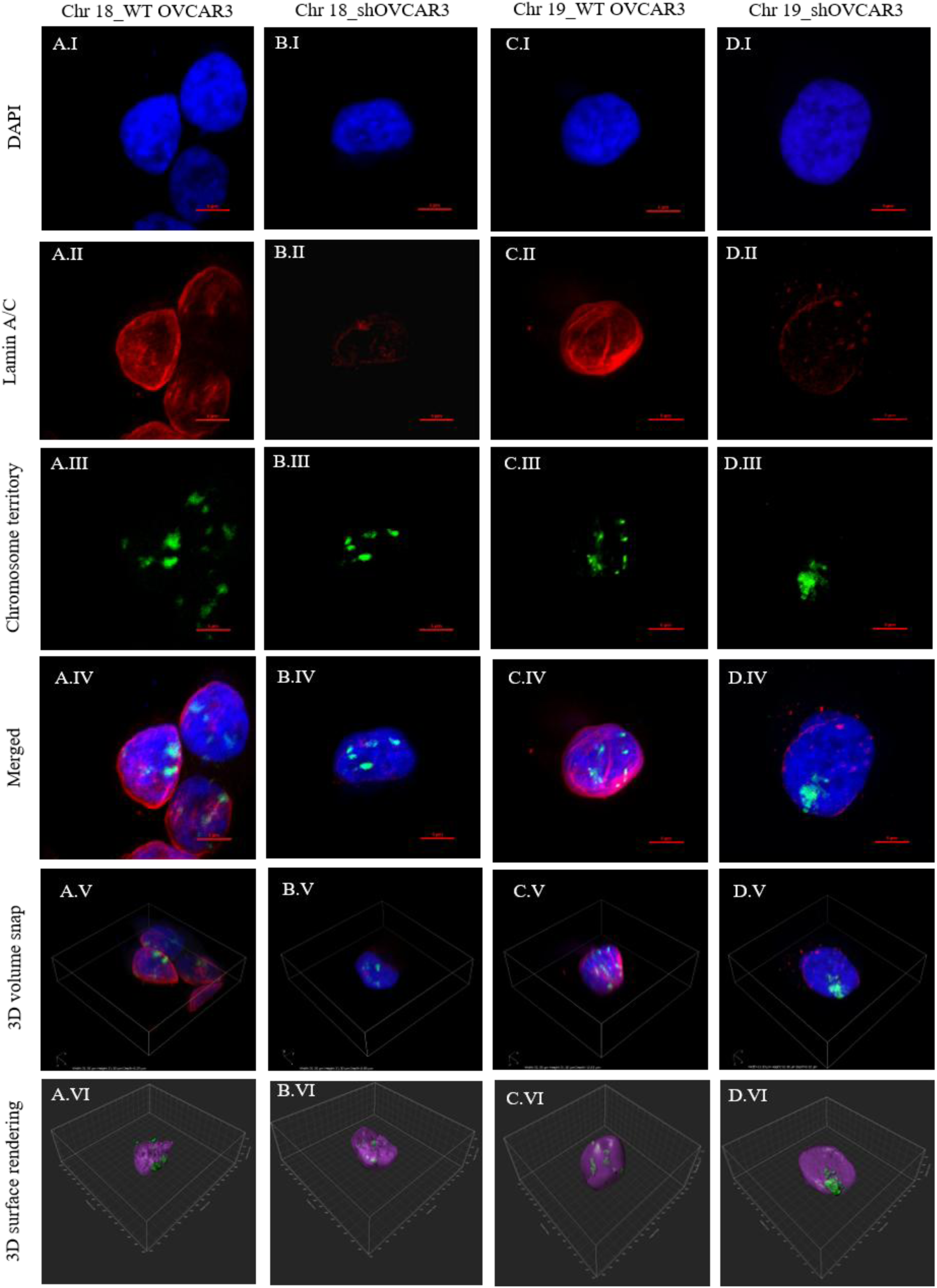
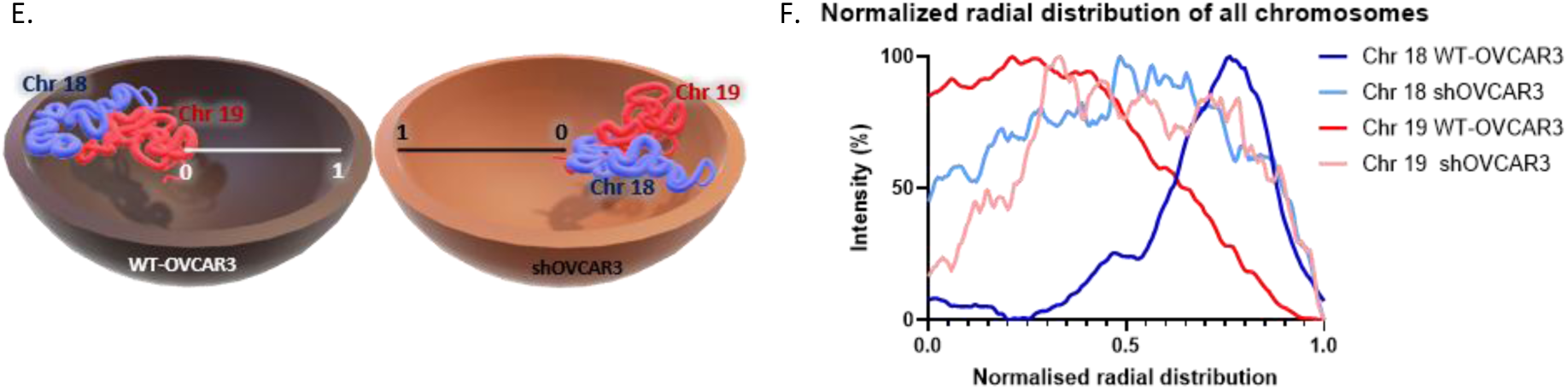
Spatial reorganization of of chromosome 18 and 19 territories in WT-OVCAR3 and shOVCAR3 nucleus. Representative confocal images (scale bar: 5 μm) from FISH and immunofluorescence experiments showing the distribution chromosome territories in: **(A)** Chromosome 18 in wild-type cells (Chr 18_WT OVCAR3), **(B)** Chromosome 18 in lamin A/C knockdown cells (Chr 18_shOVCAR3), **(C)** Chromosome 19 in wild-type cells (Chr 19_WT OVCAR3), and **(D)** Chromosome 19 in lamin A/C knockdown cells (Chr 19_shOVCAR3). Maximum intensity projection of confocal image stack, captured in 100X magnification, in individual channels illustrates: nuclear counterstain (DAPI, blue; I), the nuclear lamina (anti-Lamin A/C, red; II), and targeted chromosome territories (chromosome-specific probes, green; III). Merged multi-channel images are shown in panel IV. Corresponding 3D nuclear reconstructions are presented as 3D volume snap (V) and 3D surface rendition images (VI). **(E)** 3D schematic representing the model adopted for 3D distance measurements of chromosome 18 (blue) and chromosome 19 (red) territories between WT-OVCAR3 and shOVCAR3 nuclei (nuclear center = 0, nuclear periphery = 1). **(F)** Normalized radial distribution curves quantifying fluorochrome intensity of chromosome 18 (blue traces) and chromosome 19 (red traces) from the nuclear center to the nuclear periphery in WT-OVCAR3 versus shOVCAR3 cells.

### Polymer model validation of differential chromosome movement in lamin A/C knockdown scenario

As an orthogonal approach to validate our experimental observation pertaining to relocalisation of Chr 18 & 19, as a complement, we performed a coarse-grained polymer simulation with a far excess of lamin B particles over lamin A, which was knocked down. Starting from homogeneous initial conditions, the simulations exhibited spontaneous compartmentalisation of heterochromatin driven by differential interaction strengths. Heterochromatin beads progressively merged into dense, spatially compact clusters, while euchromatin remained more dispersed along the chain. Lamin A and B particles independently accumulated at the simulated nuclear periphery via the confining wall potential, forming a stable surface layer largely separated from the chromatin polymer. This behaviour reflected block-segregation-driven domain formation along a single covalently connected chromatin chain, arising from the differential pairwise affinities, rather than condensation of independently diffusing molecular species as in classical liquid–liquid phase separation. We therefore proceeded to describe the resulting heterochromatin clusters as compartmentalised domains rather than phase-separated liquid droplets, consistent with the single-chain architecture of the model. In both wild-type and lamin-A-knockdown conditions, progressive peripheral accumulation of lamin A and B particles (Fig. 3) is consistent with the asymmetric wall potential governing lamin-wall interactions (71), and was observed independently of direct lamin–lamin attraction. Surface accumulation of lamins, by contrast, followed a more consistent temporal pattern. In other words, lamin A and B particles in both the wild-type and knockdown state were already substantially enriched at the peripheral wall by the earliest sampled timestep (step 10,000) and remained concentrated there throughout the end of the simulation (4 × 10^5^steps), indicating that peripheral lamin localisation was established early and maintained throughout, in contrast to the more gradual and fluctuating evolution of heterochromatin cluster positioning. Collectively, these results demonstrate that a minimal coarse-grained model incorporating differential lamin interactions and a confining nuclear-envelope-like potential is sufficient to reproduce two key organisational features observed in nuclear chromatin: namely, the lamin accumulation at the nuclear periphery, and heterochromatin compartmentalisation that is modulated by lamin A/B composition.

**Fig. 3.**
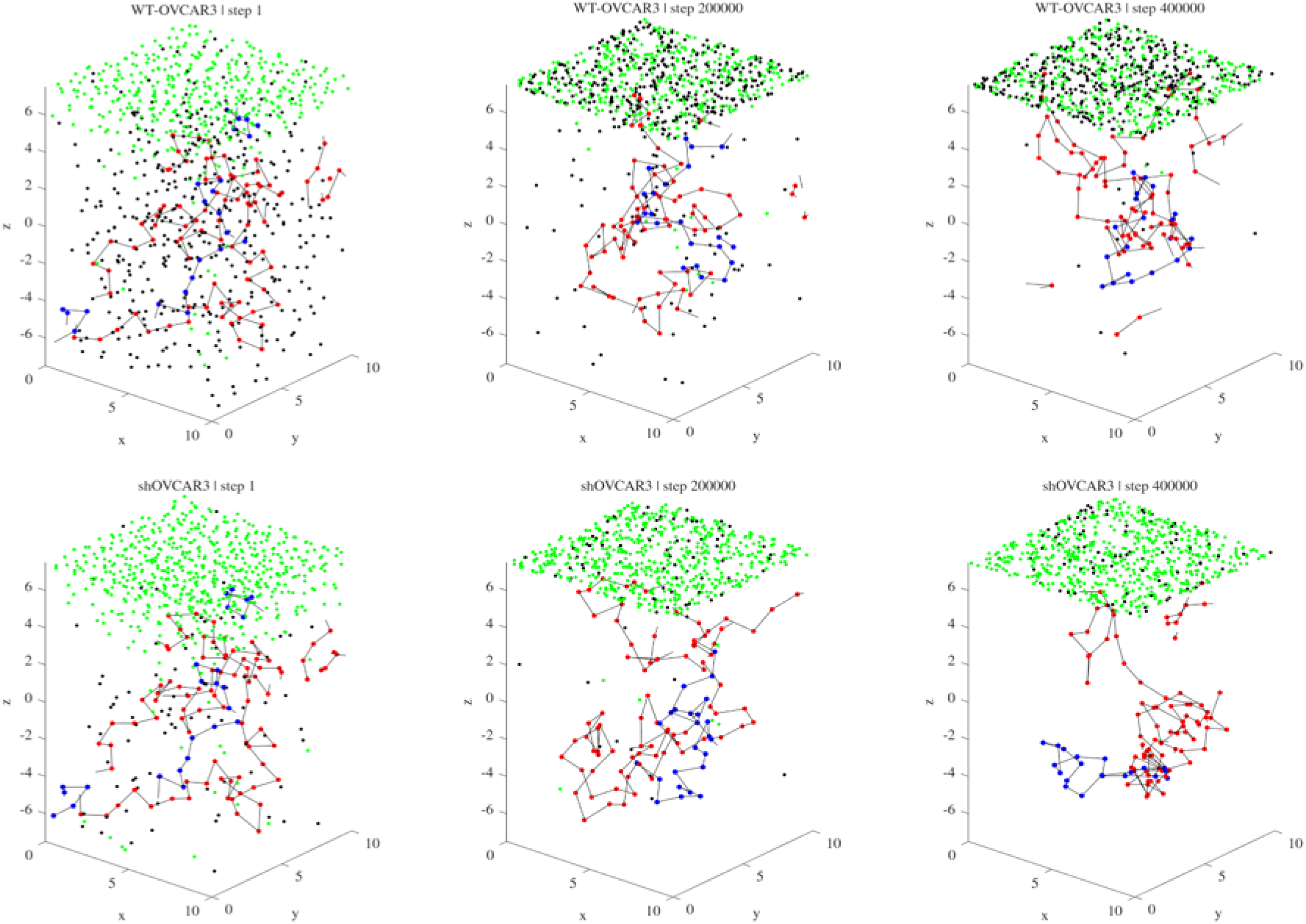
Representative simulation snapshots of chromatin and lamin organization in WT-OVCAR3 and shOVCAR3 scenario. Snapshots for WT-OVCAR3 (top row) and shOVCAR3 (bottom row) showcasing simulation steps 1, 200,000, and 400,000 (from left to right). Colors represent euchromatin (red), heterochromatin (blue), lamin A (black), and lamin B (green). Spatial axes (x, y, z) are expressed in units of the bead diameter *σ*.

We calculated the distance-resolved distributions of euchromatin (EC) and heterochromatin (HC) as a function of their distance from the confining peripheral surface. The resulting distributions, P(z), were evaluated throughout the simulation and compared between the initial configuration (step 1) and the final configuration (step 400,000) for WT-OVCAR3 and shOVCAR3 conditions (Fig. 3). The envelopes represented by thin blue lines demonstrate the distributions at intermediate simulation times, while the black and blue solid lines denote the initial and final states, respectively. Compared with the initial configuration, the heterochromatin distribution underwent an evident spatial rearrangement (Fig. 4A, B), with the final distribution shifting towards the centre from the nuclear periphery in shOVCAR3 with respect to WT-OVCAR3. At the initial time, HC showed substantial probability at relatively small distances from the peripheral surface, consistent with peripheral localisation. But during the course of the simulation, this distribution progressively shifted toward larger distances. Thus, shOVCAR3 was associated with a pronounced redistribution of heterochromatin from the nuclear periphery toward the interior of the simulated nuclear volume. In contrast, no significant redistribution of euchromatin was visualised except for some temporal broadening and shifts in the position of the distribution in the lamin A/C depleted scenario when compared with wild type. (Fig. 4C, D).

**Fig. 4.**
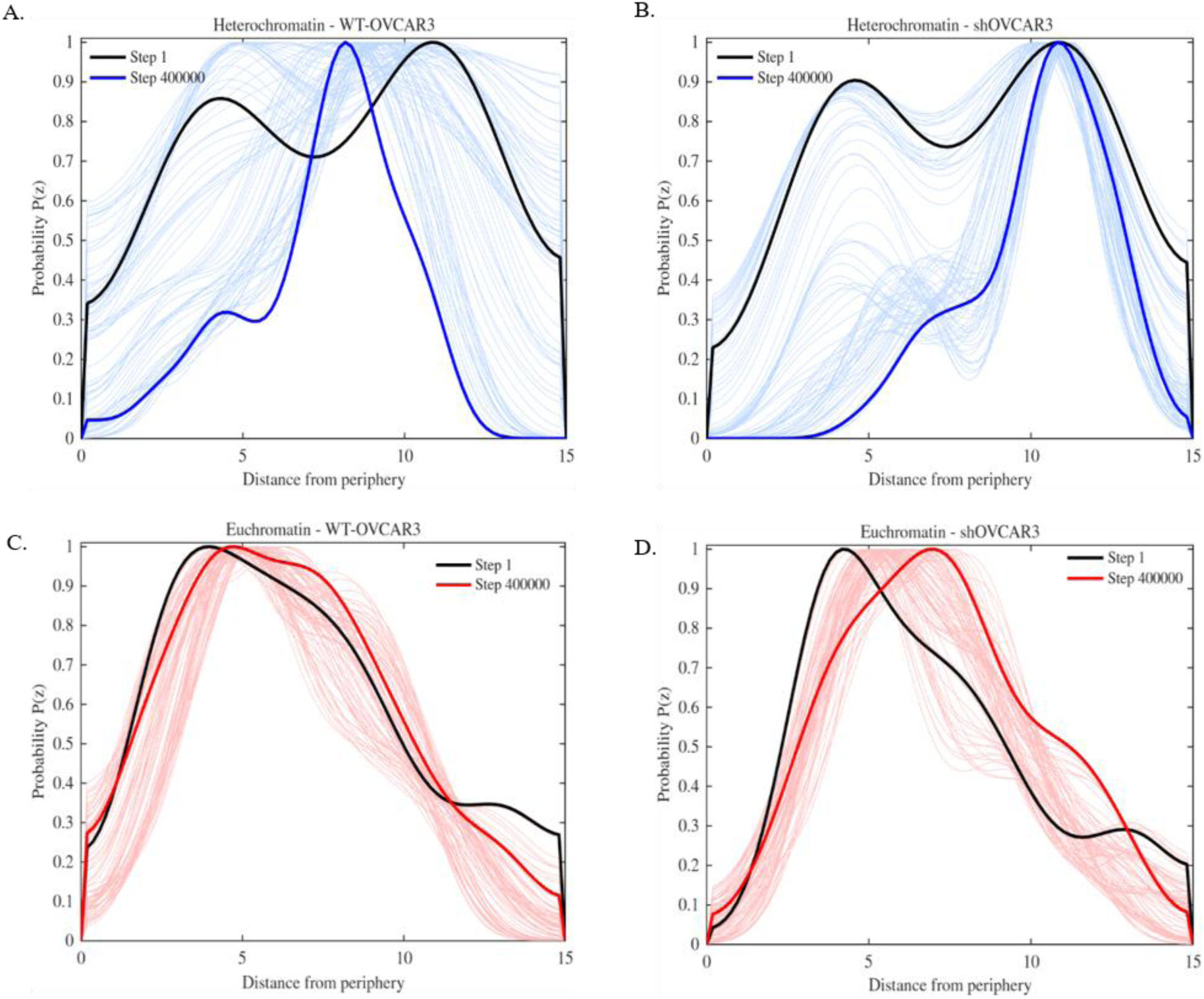
Temporal dynamics of heterochromatin organization in WT-OVCAR3 and shOVCAR3 cells. Radial probability distributions P(z) of Heterochromatin (Blue; **A, B**), Euchromatin (Red; **C, D**) as a function of distance from the nuclear periphery for WT-OVCAR3 (A,C) and shOVCAR3 (B,D). The solid black lines represent the initial configuration (step 1), while the solid coloured lines represent the final configuration (step 400,000). Thin lines show the distributions at intermediate simulation steps, illustrating the progressive redistribution of chromatin within the nucleus. Distance from the periphery is expressed in units of σ.

Together, these distance-resolved distributions complemented the cluster-size and order-parameter analyses presented above. While the clustering analysis demonstrated enhanced heterochromatin self-association in shOVCAR3 (Sup Fig.1), the peripheral-distance analysis showed that this increased clustering was accompanied by a substantial change in the spatial positioning of heterochromatin. Notably, this redistribution was selective for HC, whereas the overall EC distribution remained comparatively stable. These results further predict that reducing lamin A/C modulates the spatial relationship of HC with the nuclear periphery along with heterochromatin clustering, which corroborates the biophysical modelling by Chiang et al. (97), which also demonstrated altered physical boundary conditions of heterochromatin in a progeroid model. In summary, the Coarse-grained (CG) simulation led us to the same conclusion that emerged from 3D-FISH data.

### Spatial evolution of interchromosomal contacts and compartment switching with lamin A/C expression

With the foundational insights into chromosome reorganisation obtained from 3D FISH and CG simulations, we set forth the aim to delineate the implications at the level of inter/intra chromosomal contacts and the corresponding TADs, if any, via high-throughput chromosome conformation capture (Hi-C) performed in WT and shOVCAR3 cells. We first assessed large-scale chromosome positioning by analysing inter-chromosomal interactions. The Hi-C contact matrices for all chromosomes showed a visible increase in inter-chromosomal contact frequencies in lamin A knockdown cells compared to wild-type (Fig. 5A). To quantify this, we calculated the inter-chromosomal interaction ratio for each chromosome, defined as the proportion of its interactions occurring with other chromosomes versus the overall interactions. This analysis revealed a highly significant increase in inter-chromosomal interactions in shOVCAR3 cells (p=7.7E-11) (Fig. 5B), indicating that lamin A is required for the proper segregation of chromosome territories. Subsequently, we asked whether lamin A/C depletion affects the organisation of A and B compartments of each chromosome, which in turn are associated with active (A) and inactive (B) chromatin states. Genome-wide analysis of all autosomal chromosomes identified 1,233 stable A-compartment bins and 1,136 stable B-compartment bins, compared with 143 B-to-A and only 126 A-to-B transitions, demonstrating that lamin depletion primarily induces localised compartment remodelling rather than extensive chromosome-scale reorganisation (Fig. 5C). Bins assigned to the A compartment consistently contained a higher number of annotated genes than bins assigned to the B compartment. Genome-wide, stable A-compartment bins contained an average of 31.6 genes per 1 Mb bin, whereas stable B compartments possessed 23.2 genes per bin. Similar trends were observed for chromosome-specific analyses, with chromosome 18 (Fig. 5D), which is reported to be gene-poor, containing an average of 24.7 genes per A-compartment bin versus 20.3 genes per B-compartment bin, while chromosome 19 (Fig. 5E), which is gene-rich, exhibited substantially higher overall gene density (65.5 versus 57.4 genes per bin, respectively). The interaction heatmap from the differential Hi-C matrix (log_2_(shOVCAR3/WT-OVCAR3)) showed widespread gains in long-range contacts while preserving short-range organisation along the neutral diagonal. For chromosome 18, the majority of genomic bins retained their compartment identity following lamin knockdown. Among the 73 informative 1 Mb bins, 19 remained in the A compartment, and 45 remained in the B compartment, whereas 8 bins underwent A-to-B transitions and only 1 bin exhibited a B-to-A transition. These compartment toggling events were primarily localised to the 26–27 Mb, 34–36 Mb and 58–64 Mb regions (Fig. 5F). Chromosome 19 exhibited a greater degree of localised remodelling despite maintaining an overall stable compartment architecture. Among 55 informative bins, 20 remained in the A compartment, and 26 remained in the B compartment, while 5 bins underwent B-to-A transitions and 4 bins underwent A-to-B transitions. The strongest B-to-A transitions occurred within the 40–41 Mb and 53–55 Mb regions, whereas the most pronounced A-to-B transition occurred near 18–19 Mb (Fig. 5G). From quantification of intercompartment switching, a preponderance of transition towards the A compartment was found, which led us to reason that lowering the level of lamin A/C might lead to an increase in transcriptional activity. These findings indicate that the peripheral and nucleoplasmic distribution of lamin A/C plays an important role in delineating A and B compartments in the genome, interacting with both euchromatin and heterochromatin regions, as evident from 3D FISH. Also, the expression level of A-type lamins plays a deterministic role in the fate of intercompartmental switching, essentially modulating the stability of compartment identity and the spatial segregation between compartment types.

**Fig. 5.**
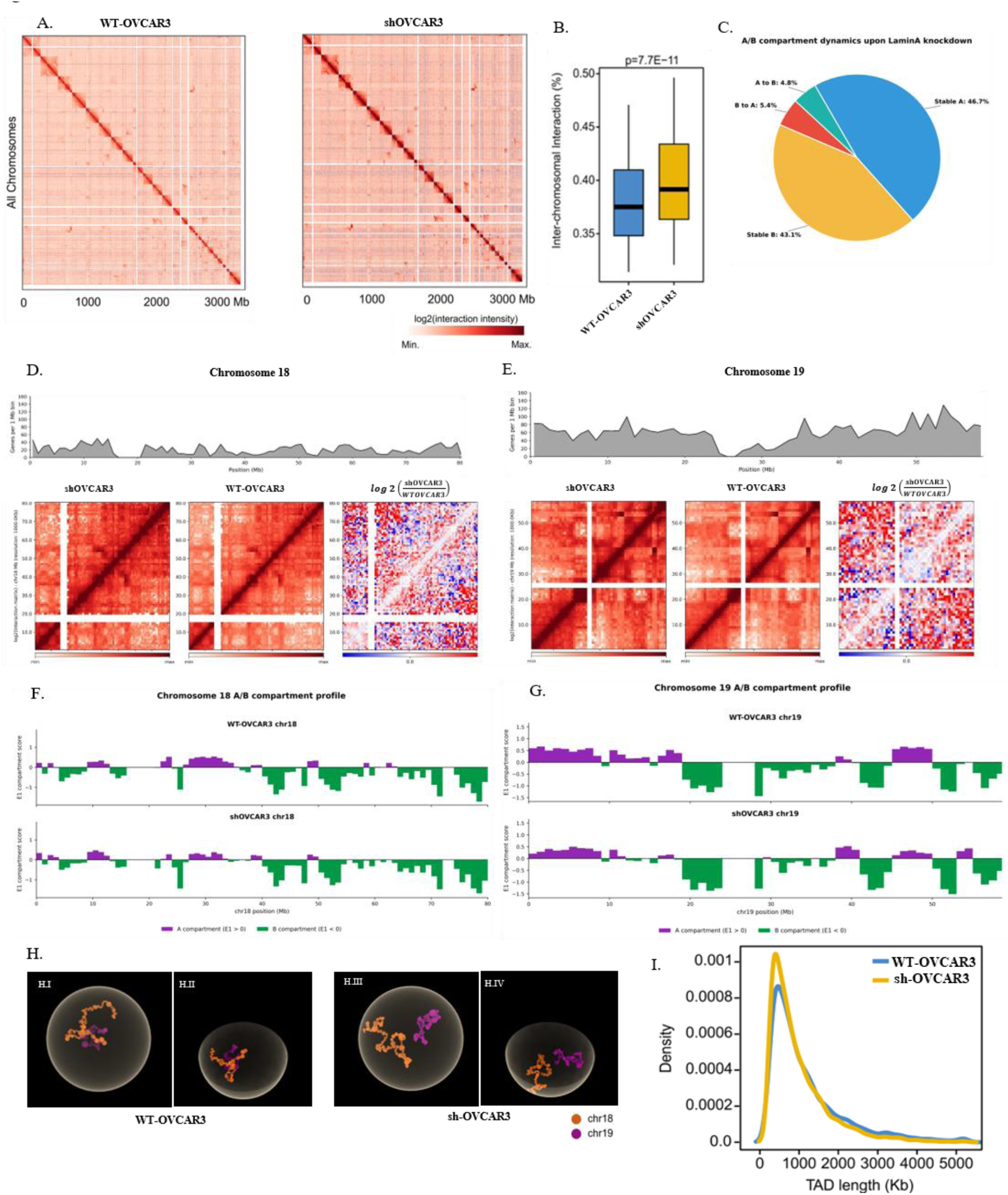
Lamin A/C knockdown alters genome-wide chromatin architecture and spatial organization. **(A)** Whole genome Hi-C contact matrices at 500 kb resolution for WT-OVCAR3 (left) and shOVCAR3 (right), showing normalized log2 interaction intensities. **(B)** Boxplot comparing the inter-chromosomal interactions ratio of WT-OVCAR3 versus shOVCAR3 cells. The horizontal line shows median values whereas the box shows the inter-quartile range. **(C)** Pie chart showing genome-wide A/B compartment dynamics upon Lamin A/C knockdown, colors representing proportions of Stable A (blue), Stable B (yellow), B-to-A transitions (red), and A-to-B transitions (green). **(D, E)** Gene density profiles per 1 Mb bin (top) and high-resolution (100 kb) Hi-C interaction heatmaps (bottom left) with corresponding differential interaction matrices (log₂(shOVCAR3/WT-OVCAR3); bottom right) for Chromosome 18 (D) and Chromosome 19 (E). **(F, G)** First principal component (E1) profiles for WT-OVCAR3 (top tracks) and shOVCAR3 (bottom tracks), The purple color represents active compartment A, whereas green color represents inactive compartment B along Chromosome 18 (F) and Chromosome 19 (G). **(H)** Representative 3D genome reconstruction illustrating the nuclear localization of Chromosome 18 (orange) and Chromosome 19 (purple) in WT-OVCAR3 (H. I, H. II) and shOVCAR3 (H.III, H.IV) nuclei, I and III displaying standard front-facing view while II and IV showing angled perspective view of the same model. **(I)** Line plot representing density distribution of Topologically Associating Domain (TAD) lengths in WT (blue) versus shOVCAR3 (yellow) cells.

To revisit the pattern of spatial distribution from Hi-C contact matrices, 3D genome reconstructions of WT and sh conditions were modelled using Chrom3D. Although we could see an apparent partial detethering of chromosome 18 from the periphery with a redistribution radially towards the centre in the knockdown condition in the unclipped 3D nuclear volumes as well as angled perspectives (Fig. 5H), we acknowledge that models from Chrom3D are only predictive or illustrative representations, acting only as qualitative models of chromosomal topology.

In contrast to the more pronounced effects on compartments and chromosome territories, the depletion of lamin A/C had a lesser impact on topologically associating domains (TADs), which are kilobase structural units of chromatin. The distribution of TAD lengths was nearly identical between WT and shOVCAR3 cells (Fig. 5I). This result conforms to the lamin B1 knock-out study where practically no change was observed in the distribution of TAD lengths. Nonetheless, we found an overall increase in the number of TADs in the lamin A/C knockdown scenario, indicating newly formed TADs upon depletion of the nucleoplasmic scaffold (Supplementary file 1, sheet TAD count). Taken together, our results demonstrate that lamin A depletion disrupts chromosome territory segregation, significantly increases inter-chromosomal contact frequencies, and modulates compartment identity, while preserving the local TAD structure of the genome. This brings us to a juncture where we now interrogate how the lamin A/C mediated perturbations in inter-chromosomal contacts affect the formation of transcription hubs driving a plethora of genes involved in oncogenic transformation.

### Rewiring of oncogenic super enhancer–promoter contacts

To determine whether lamin A/C depletion affects the higher-order chromatin organisation of previously characterized OVCAR3 super-enhancers (SEs), 86 published OVCAR3 SE loci (67) were integrated with the A/B compartment profiles of WT and shOVCAR3 cells. The majority of SEs retained their compartment identity, with a smaller subset exhibiting compartment switching as a downstream effect of the knockdown. 55 of 86 loci (64.0%) remained in the A compartment and 23 (26.7%) in the B compartment. In contrast, five SEs (5.8%) transitioned from the B to the A compartment, whereas only one SE (1.2%) showed an A-to-B transition. Two loci could not be assigned in both conditions (Fig. 6A). Analysis of the corresponding compartment eigenvector scores identified SE56, SE86 and SE36 as the strongest B-to-A switching loci. SE56 exhibited the largest change, from an E1 value of −0.355 in WT cells to +0.293 following lamin knockdown (ΔE1 = +0.648), followed by SE86 (−0.399 to +0.175; ΔE1 = +0.574) and SE36 (−0.187 to +0.276; ΔE1 = +0.463. Two additional loci, SE20 and SE77, were classified as B-to-A transitions but contained E1 values close to the compartment boundary in one condition. Similarly, the only A-to-B event, SE95, shifted from +0.223 to −0.010 and therefore represented a comparatively weak transition close to the compartment boundary (Fig. 6B).

**Fig. 6.**
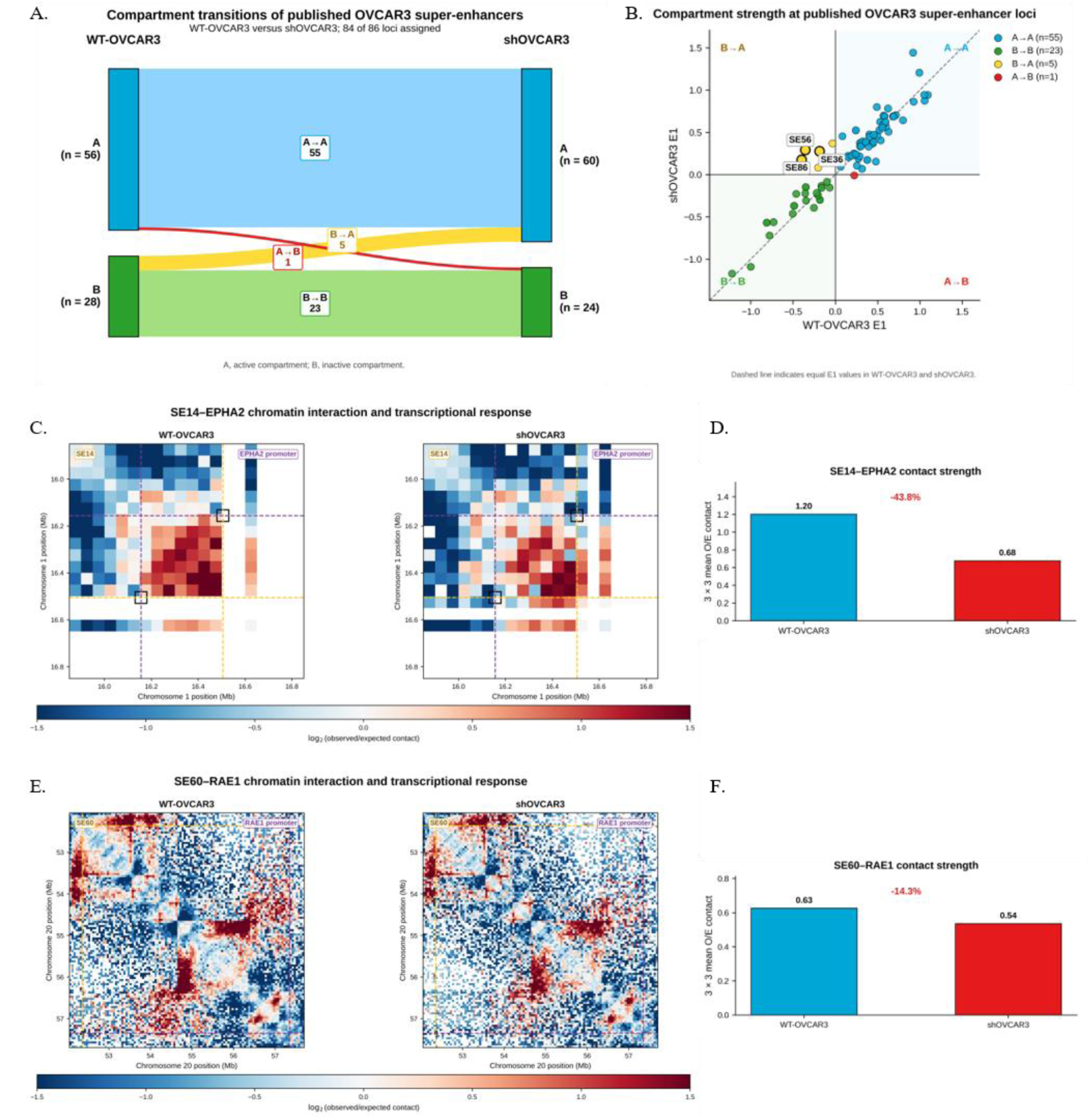
Overexpressed lamin A/C maintains A/B compartmentalization and chromatin interactions at oncogenically implicated super-enhancer loci in WT-OVCAR3 cells. (A) Alluvial representation of A/B compartment transitions across 84 assigned super-enhancer (SE) loci between WT-OVCAR3 and shOVCAR3 cells. (B) Scatter plot comparing eigenvector 1 (E1) compartment scores at SE loci between WT-OVCAR3 and shOVCAR3 cells. The dashed diagonal indicates equal E1 values in both conditions. Loci in the upper-left quadrant represent B-to-A transitions, whereas loci in the lower-right quadrant represent A-to-B transitions. SEs showing the strongest B-to-A transitions are highlighted. (C) Balanced observed/expected HiC contact maps at 50-kb resolution featuring the SE14–EPHA2 locus in WT-OVCAR3 (left) and shOVCAR3 (right) cells. Dashed orange and purple lines indicate the SE14 and EPHA2-promoter bins, respectively, and black boxes mark the corresponding enhancer– promoter contact pixels. Color intensity in both WT and sh-conditions scales log₂(observed/expected contact). (D) Quantification of the mean observed/expected contact within a 3 × 3 bin window centered on the SE14-EPHA2 promoter bins in WT-OVCAR3 and shOVCAR3 cells. (E) Balanced observed/expected Hi-C contact maps covering SE60–RAE1 locus in WT-OVCAR3 (left) and shOVCAR3 (right) cells. The SE60 and RAE1-promoter positions are indicated by dashed orange and purple lines respectively. Color intensity represents log₂(observed/expected contact). (F) Quantification of the mean observed/expected contact within a 3 × 3 bin window centered on the SE60 and RAE1-promoter bins, comparing contact strength between WT-OVCAR3 and shOVCAR3 cells

At this point, we justify the rationale for revisiting the previously reported oncogenically implicated SE-promoter interactions in OVCAR3 because we intended to determine whether depletion of lamin A/C induced any further perturbations in their 3D chromosome contacts. At the SE14–EPHA2 locus, normalised Hi-C analysis showed a reduction in local SE–promoter interaction following lamin A/C knockdown (Fig. 6C). The mean observed/expected contact strength decreased from 1.20 in WT-OVCAR3 to 0.68 in shOVCAR3 (Fig. 6D), corresponding to an approximately 43.8% reduction in SE14–EPHA2 contact. Similarly, at the SE60–RAE1 locus, the normalised Hi-C interaction also decreased following the knockdown (Fig. 6E), with the mean observed/expected interaction decreased from 0.63 in WT-OVCAR3 to 0.54 in shOVCAR3, which corresponded to an approximate 14.3% reduction (Fig. 6F).

### Lamin A/C depletion induced abrogation of oncogenic proliferation

So far, we have established a global reorganisation of chromosome territories and repurposing between A and B compartments, along with an altered SE-promoter interaction following lamin A/C knockdown. Along this line of investigation, the next relevant question was how the transcriptome was affected as a downstream phenomenon. Therefore, we resorted to RNA Sequencing analysis which at the foremost confirmed an efficient depletion of the targeted transcript, LMNA appearing amongst the most strongly downregulated genes with a log₂ fold change of −1.91 in shOVCAR3 relative to WT-OVCAR3 (Fig. 7A). Using the differential expression threshold of an absolute log_2_fold-change ≥ 0.585 (fold change ∼ 1.5x) and adjusted value < 0.05, a total of 1833 genes were found to be differentially expressed comprising 1334 upregulated and 499 downregulated genes in shOVCAR3 cells. Principal component analysis captured 83.5% total variance among the samples of shOVCAR3 and WT-OVCAR3, showing clear separation (Fig. 7B). Consistent with the separation, the hierarchical clustering of significant differentially expressed genes segregated the samples into distinct shOVCAR3 and WT-OVCAR3 clusters (Fig. 7C). The 50 top differentially expressed genes are shown in Fig. 7D where LMNA appeared among the prominent genes distinguishing wild-type and knockdown cells. The lists of all significant differentially regulated genes are provided in - DE_up/downregulated_in_SH_1.5x sheets in the supplementary file 1.

**Fig. 7.**
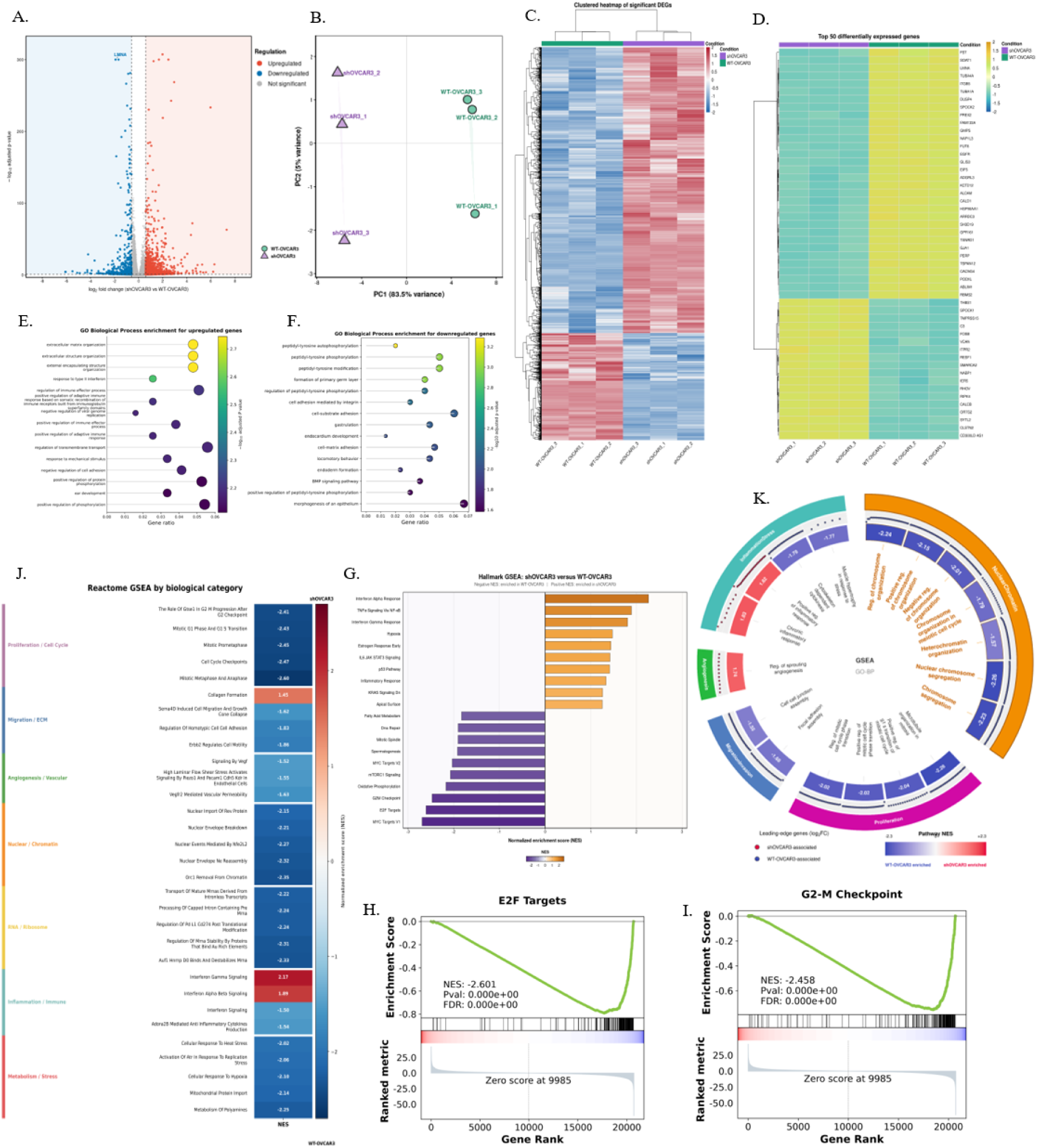
Transcriptomic and functional consequences of lamin A/C knockdown in OVCAR3 cells. **(A)** Volcano plot of differentially expressed genes (DEGs) between shOVCAR3 and WT-OVCAR3 control with n = 3 biological replicates per group. Significantly upregulated genes with log₂ fold-change ≥ 0.585, are highlighted in red, and downregulated genes with log₂ fold-change ≤ 0.585 in blue. Non-significant genes are marked in grey. LMNA, the targeted lamin gene, is labeled. **(B)** Principal component analysis of variance-stabilized RNA-seq data from shOVCAR3 and WT-OVCAR3 replicates. **(C)** Hierarchical clustering of DEGs across individual biological replicates. The color scale represents relative expression ranging from high (red) to low (blue). **(D)** Heatmap of the top 50 differentially expressed genes. displaying relative expression patterns with color scale. **(E, F)** Gene Ontology (GO) Biological Process enrichment analysis of upregulated (E) and downregulated (F) genes, in shOVCAR3 with respect to WT-OVCAR3. Circle position indicates gene ratio, circle size indicates the number of genes associated with each term, and colour intensity represents statistical significance. **(G)** Hallmark gene-set enrichment analysis (GSEA) summary. Bar chart displaying positive normalized enrichment scores (orange) indicate enrichment in shOVCAR3 cells, whereas negative scores (purple) indicate enrichment in WT-OVCAR3 cells. **(H,I)** Gene-set enrichment score plots for E2F Targets (H) and G2–M Checkpoint (I), respectively. Vertical black lines indicate the positions of gene-set members within the ranked gene list. **(J)** Reactome pathways identified by GSEA are shown according to their normalized enrichment score in bar plot. Significantly enriched pathways are grouped into key biological categories, indicated by the color-coded annotation bar on the left. **(K)** Representation of enriched Gene Ontology biological processes grouped into major functional categories in a circular plot showcasing pathway normalized enrichment scores (NES), where individual points represent leading-edge genes.

GO analysis revealed upregulation of specific functional signatures associated with extracellular matrix organisation, extracellular structure organisation, and external encapsulating structure organisation in shOVCAR3 (adjusted P = 0.0018 for each), thereby indicating substantial remodelling of extracellular structural pathway associated transcripts (Fig. 7E). On the other hand, significantly downregulated genes were enriched for processes associated with regulation and phosphorylation of peptidyl-tyrosine, integrin-mediated adhesion, cell-substrate and cell-matrix adhesion (Fig. 7F). Therefore, lamin A/C knockdown leads to perturbations in extracellular matrix as well as alterations in signalling and cellular adhesion pathways. GSEA revealed, without applying a defined fold change, broad suppression of cell-cycle, chromatin-associated programmes and stress-response transcriptional programmes following lamin A/C knockdown. Hallmark GSEA pathways (Fig. 7G) supported a similar phenotype as Reactome pathways. Among the most negatively enriched gene sets were MYC Targets V1, G2-M Checkpoint, E2F Targets, mTORC1 Signalling and Mitotic Spindle, all of which belonged to 11 significant hallmarks that were previously identified from GSEA analysis, having a strong positive correlation with ovarian cancer (114). The E2F Targets (Fig. 7H) and G2-M Checkpoint (Fig. 7I) showed strong negative enrichment of NES = −2.601 and -2.458 respectively, with an FDR < 0.001, which, along with several other pathways from Hallmark GSEA analysis (Supplementary file 1, sheet GSEA_Hallmark_SH_vs_WT), are indicative of cell cycle arrest and delayed cellular proliferation. The GSEA Reactome (Fig. 7J) analysis likewise elucidated consistent negative enrichment of pathways involved in cell-cycle progression, mitosis, DNA replication and chromosome dynamics, indicating that genes belonging to such pathways are preferentially expressed in WT-OVCAR3-associated end of the ranked transcriptome. The negatively enriched pathways included several ones directly related to nuclear and chromatin organisation, including nuclear envelope breakdown (NES = −2.21), Nuclear events mediated by NFE2L2 (NES = −2.27), Nuclear envelope reassembly (NES = −2.32), ORC1 removal from chromatin (NES = −2.35).Mitotic Metaphase and Anaphase (NES= -2.60), Separation of Sister Chromatids (NES = −2.58), Resolution of Sister Chromatid Cohesion (NES = −2.49), Cell Cycle Checkpoints (NES = −2.47), Mitotic Prometaphase (NES = −2.45), S Phase (NES = −2.43), Mitotic G1 Phase and G1/S Transition (NES = −2.43), Mitotic G2/G2-M Phases (NES = −2.40) and G2/M Checkpoints (NES = −2.39). DNA replication was similarly negatively enriched (NES = −2.32) (supplementary file 1, sheet GSEA_Reactome_SH_vs_WT). Thus, these findings clearly demonstrated the correlation between LMNA depletion and concomitant transcriptional changes in nuclear-envelope, chromatin and chromosome-associated programmes, consistent with the fact that lamin A/C moonlights in diverse roles to maintain nuclear architecture and homeostasis. The findings from GSEA Reactome analysis cohere with the insights from GO-GSEA analysis (supplementary file 1, sheet GSEA_GO_BP_SH_vs_WT). Interestingly, heterochromatin organisation (Fig. 7K) showed a negative enrichment (NES = −1.57), indicating preferential representation of genes contributing to heterochromatin organisation towards the WT-OVCAR3-associated end of the ranked transcriptome. It can be aptly concluded here that insights from GO-GSEA analysis are largely in line with our explanation from 3D-FISH and CG simulations. In summary, these results indicated suppression of proliferative programmes, broad downregulation of regulatory networks governing structural chromosome integrity, deregulation of chromosome organisation, as well as specialised heterochromatin organisation following LMNA knockdown.

Strikingly, lamin A/C depletion was found to attenuate ERBB/EGFR signalling in shOVCAR3, with EGFR or ERBB1 being amongst the top 50 differentially downregulated genes (Fig. 7D). Reactome GSEA showed a coordinated reduction of multiple ERBB-associated signalling pathways following LMNA depletion (supplementary file 2). All 10 ERBB-related Reactome pathways showed negative NES values, of which 7 exhibited significant preferential enrichment toward WT-OVCAR3. We identified core ERBB-associated genes which led to these multiple pathway enrichments (Fig. 8). A prominent core formed through the intersection of 10 pathways consisted of EGFR, ERBB4, NRG1, BTC and NRG2. EGFR, ERBB4, NRG1, and BTC each contributed to 9 of the 10 ERBB-related pathways, while NRG2 occurred in 8 pathways. These genes thus represent a recurrent receptor–ligand module underlying much of the ERBB-associated GSEA signal. The downstream RAS signalling components KRAS and HRAS each contributed to six ERBB-related pathways, while HSP90AA1 and PTPN12 occurred in four. PLCG1 contributed to three pathways. The three AKT paralogues, AKT1, AKT2 and AKT3, were represented in both- Signaling by ERBB2 pathway and downregulation of ERBB2 Signaling. The motility-associated module contained MEMO1, RHOA and DIAPH1, which also featured in both the broad ERBB2 signalling pathway and the more specific ERBB2 Regulates Cell Motility pathway. The latter was one of the most strongly and significantly WT-associated ERBB pathways (NES = −1.86, FDR = 0.0012) (Fig. 7J). STRING analysis of genes connected to ERBB-associated signalling pathways indicated EGFR to be positioned within the densely interconnected central region with a notable contribution to nine of the ten ERBB-related pathways. EGFR expression across ovarian cancer subtypes was reported earlier to be a driver of rapid tumour cell proliferation, higher grade, and reduced progression-free survival (115, 116). The association of EGFR expression with endogenously overexpressed lamin A/C intrigued us to investigate whether a functional or physical link exists between these two proteins. Excavating public interactome databases (BioGRID) revealed a direct association of lamin A/C with EGFR in experimental protein–protein interaction datasets. In the study by Liccardi et al. (117) characterising DNA repair complexes involving ERCC1, LMNA was found in the baseline interactome of nuclear EGFR in high-throughput affinity capture-mass spectrometry screens. Later, Dittmann et al. (118) confirmed that nuclear-translocated EGFR co-localises with LMNA-containing ribonucleoprotein complexes via proteomic screening, co-immunoprecipitation and confocal microscopy experiments. The observation from Dittmann et al. reporting that EGFR physically interacts with lamin A/C to regulate radiation-induced stress responses, along with other reports uncovering the role of elevated lamin A/C as the major driver in double-strand break repair pathways (HR and NHEJ) in aggressive HGSOC (39), indicates that lamin A/C driven DNA repair pathways mirror EGFR-driven survival mechanisms. Lamin A/C depletion leading to reduced EGFR levels highlights a critical regulatory dependency, suggesting that endogenous lamin A/C overexpression in HGSOC is required to maintain normal EGFR homeostasis and downstream oncogenic signalling.

**Fig. 8.**
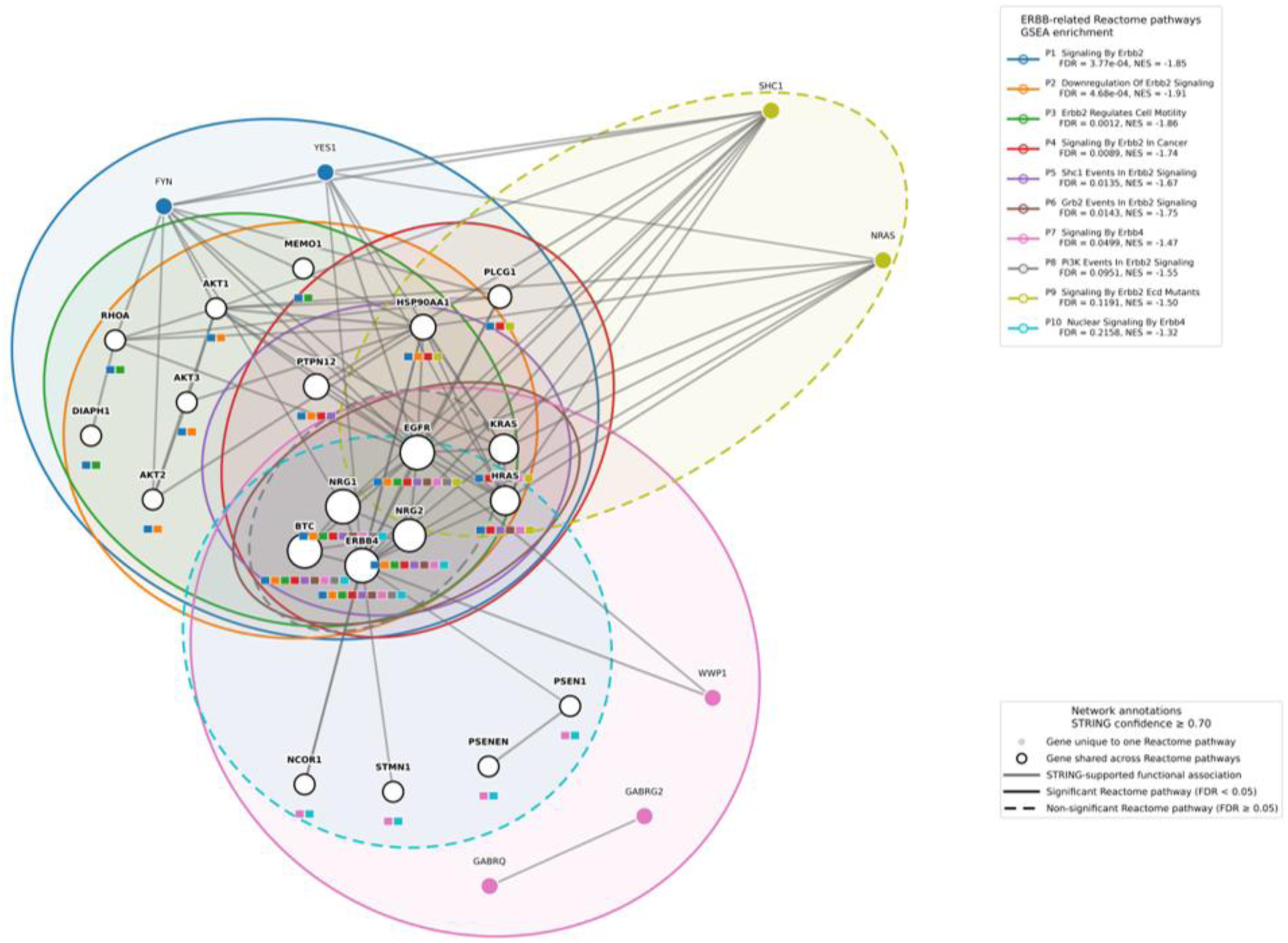
Functional crosstalk and interaction network of genes contributing to ERBB-related Reactome pathways. High-confidence STRING interaction network with interaction score ≥0.70 (grey edges), of genes enriched in ERBB-related Reactome pathways (P1–P10) identified by GSEA. Gene nodes with open circles denote genes shared across multiple Reactome pathways, and filled circles indicate genes unique to a single pathway. Colored squares beneath each node reflect its specific pathway memberships corresponding to the color-coded legend summarized in the figure. The colored pathway regions represent GSEA-derived Reactome membership, whereas the grey connecting edges represent STRING-supported functional associations. Edge width was scaled according to STRING confidence.

The attenuation at the membrane reception level by steep downregulation of EGFR affects the immediate downstream cascade – the classic MAPK axis and PI3K/AKT. We found significant downregulation of AKT3, along with modest downregulation of AKT 1 and AKT2 (supplementary file 1, sheet DE_all_results_SH_vs_WT). PREX2, which is a classical effector in PI3K signalling (119) was listed amongst the top 50 downregulated genes. Marked upregulation of DUSP5 upon knockdown of lamin A/C silences MAP kinases by dephosphorylation (120), along with an upregulation of Spry4, a negative feedback inhibitor of the RTK signalling pathway. Spry4 has been documented widely to interfere with cell proliferation and migration (121). Indication towards aberrant proliferation was strengthened by the robust upregulation of CDKN1A (p21), a Cdk kinase inhibitor (122) and a downregulation of CCNE1 (cyclin E1), which serves as a predictive biomarker in ovarian and endometrial carcinomas (123). Last but not least, SOX 17 (SRY-box transcription factor 17), which is a highly sensitive and specific marker for ovarian and endometrial carcinomas (124), was significantly downregulated.

Taking cues from the Hi-C, the three strongest B-to-A transitions in SE56, SE86 and SE36 were then examined for association with transcriptional changes in their surrounding genomic regions. Differentially expressed genes were defined using an adjusted P value <0.05 and an absolute log₂ fold change ≥ 1.0. Within ±1 Mb of SE56, one gene fulfilled these criteria, SPRY4 (log₂FC = +1.407, adjusted P = 1.17 × 10⁻²¹), which was upregulated following lamin A/C knockdown, showing a clear concordance between a B-to-A compartment transition and increased expression of nearby genes. The transcriptional status of other B-to-A switching loci was less discernible. Within ±1 Mb of SE36, out of seven differentially regulated genes, six were downregulated and one, ASPG, was upregulated (log₂FC = +1.819, adjusted P = 3.43 × 10⁻³). At SE86, the only transcript, LOC105377727, which was strongly upregulated (log₂FC = +6.288, adjusted P = 9.44 × 10⁻¹⁶), is an uncharacterized long non-coding RNA (lncRNA). Elevated SPRY4 was found to cause depleted cell division, inducing cell-cycle arrest in several cancers (121). Masoumi-Moghaddam et al. showed significant downregulation of SPRY2 and SPRY4 in ovarian tumour tissues (125). In our study, the upregulation of SPRY4 accompanied by EGFR downregulation therefore suggests an active repression of cellular proliferation affecting RTK-driven oncogenic behaviour (126). Metabolic deprivation of asparagine upon recombinant ASPG expression was found to suppress cell proliferation in leukaemia and has been widely discussed in the context of Acute Lymphoblastic Leukaemia treatment(127, 128). Amongst very few studies in HGSOC, the work by Lorenzi et al. (129) demonstrated that human ovarian cancer cells exhibit pronounced vulnerability to enzymatic asparagine depletion. In our model, notably significant transcriptional upregulation of ASPG likely mimics the metabolic consequences of asparaginase therapy by accelerating intracellular asparagine catabolism and establishing a state of metabolic restriction, indicating a tendency towards suppressed tumour growth.

Next, we assessed the expression levels of genes exhibiting differential SE-promoter contact frequencies identified from the Hi-C data analysis. RNA sequencing analysis showed a statistically significant negative expression change in RAE1 (log₂FC = −0.212, adjusted P = 7.85 × 10⁻¹¹, (Sup. Fig.2A), consistent with the observed reduction in SE60–RAE1 chromatin interaction. Overexpressed RAE1 was found to be associated with poor overall survival in HGSOC (130). In contrast to RAE1, EPHA2 displayed a very small increase in expression (log₂FC = 0.13, adjusted P = 4.49 × 10⁻⁵) (Sup. Fig.2B) despite its reduction in contact frequency with SE14.

### Promotion of cellular proliferation of OVACR3 cells by overexpressed lamin A/C

To check the growth kinetics of OVCAR3 cells upon targeted knockdown of lamin A/C, cellular proliferation was monitored for over 120 hours in both WT-OVCAR3 and shOVCAR3 cells (Fig. 9A). Viable cell counting revealed active logarithmic growth in WT-OVCAR3 cultures between 24 hours and 72 hours post-seeding. PDT of wild-type cells calculated during this exponential phase exhibited a baseline of ∼60 hours. In contrast, shOVCAR3 cells demonstrated a pronounced growth delay across the same time window, with a PDT of ∼260 hours between 24 and 72 hours, during which time span it didn’t yet reach its exponential phase of growth. As shown in the growth curve, WT-OVCAR3 cultures displayed rapid expansion, in contrast to shOVCAR3 cells, which exhibited suppressed proliferative capacity throughout the assay timeline. Mathematical calculation of growth kinetics confirmed that lamin A/C knockdown prolonged the population doubling time significantly.

**Fig. 9.**
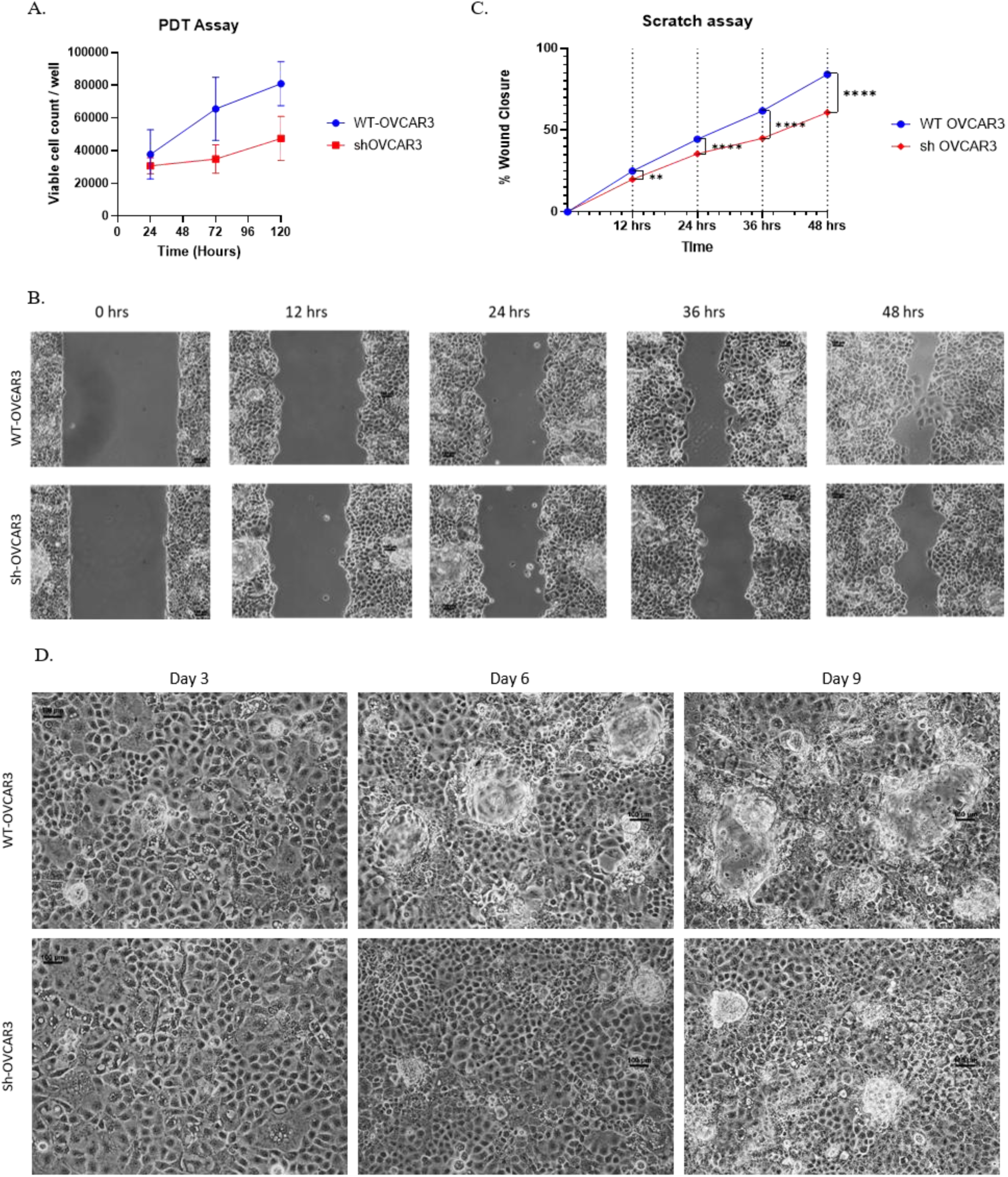
Knockdown of lamin A/C retards proliferation, migration, and results in reduced formation of spontaneous 3D spheroids from 2D monolayer in OVCAR3 cells. **(A)** Growth curve derived from population doubling time (PDT) proliferation assay showing viable cell counts per well for WT-OVCAR3 and shOVCAR3 cells measured over 120 hours. **(B)** Representative phase-contrast images captured during wound healing assay displaying cell migration into the cell-free area at 0, 12-, 24-, 36-, and 48-hours for WT-OVCAR3 and shOVCAR3 cells. (Scale bar: 100 μm). **(C)** Quantification of percentage wound closure over time corresponding to the scratch assay **(D)** Representative brightfield images capturing cell confluence and the dynamics of spontaneous formation and growth of spheroids from 2D monolayer of WT-OVCAR3 and shOVCAR3 cells. Microscopic evaluation was done for over a 9-day period post-seeding (imaged at Days 3, 6, and 9). (Scale bar: 100 μm).

Taking cues from PDT assay, we tried to understand any concurrent effect of lamin A/C deletion on migratory behaviour by performing wound-healing assay across a 48-hour time course (Fig. 9B). Beginning with uniform initial scratch widths, a statistically significant reduction in percentage wound closure emerged in shOVCAR3 cells within 12 hours and became increasingly pronounced at 24 hours and 36 hours (Fig. 9C). By the 48 hours endpoint, WT-OVCAR3 cells achieved ∼84% closure, whereas sh-OVCAR3 cells lagged significantly at ∼61% with a mean difference of 23%. Collectively, these findings demonstrate that endogenously overexpressed lamin A/C aids directional cell motility and wound closure kinetics in OVCAR3 cells.

When compared, spontaneous 3D spheroid formation propensity from 2D monolayer culture was found to be much more aggressive in WT-OVCAR3 compared to shOVCAR3. Captured images of random areas on days 3, 6, and 9 revealed progressive, visible enlargement in the number and size of budding spheroids in WT condition over time but a comparatively reduced growth and sparse assembly of budding spheroids in sh-OVCAR3 (Fig. 9D). These findings indicate that the knockdown not only attenuates cellular motility but also significantly alters its capacity for spontaneous 3D multicellular aggregation and spheroid growth dynamics.

The phenotypic impairments, including increased PDT, significant loss of cell motility and impaired in vitro wound healing, support the observations from RNA sequencing and Hi-C. The marked suppression of the Migration / ECM module in Reactome GSEA of the shOVCAR3 (Fig. 7J), with significant negative enrichment of broader ERBB signalling cascades (Fig. 8), accounts for the attenuated cellular proliferation along with broadly attenuated downstream signal propagation through the canonical MAPK and PI3K/Akt survival axes. The overall attenuation of oncogenically important SE-promoter interaction upon the diminished lamin A/C scaffold in knockdown cells also contributes to this reduced proliferation. Marked upregulation of ASPG and SPRY4, besides downregulation of RAE1, contributes to antiproliferative biochemical modulations in shOVCAR3.

ERBB2/EGFR signalling was reported to contribute to the growth of cells as spheroids, which is the primary mechanism of metastasis of ovarian cancer. Knockdown of EGFR was reported to reduce spheroid formation by more than 70% in HGSOC cells (131). EGFR or ERBB2 upregulation in nonadherent cells was not only established to increase spheroid-initiating capacity but also found to contribute to the survival and growth of nonadherent spheroids. Widespread shutdown of this ERBB signalling network in our case was highly consistent with the reduced spheroid formation in shOVCAR3.

### Lamin A/C deficiency leads to reduced tumourigenic potential of OVCAR3 cells

Having demonstrated the pro-proliferative role of endogenously overexpressed lamin A/C in OVCAR3 cells from our in-vitro experiments, we sought to delineate its role in inducing tumourigenic capacity in a physiologically relevant system. We examined comparative xenograft tumour formation and progression in nude mice injected with WT and shOVCAR3 cells, represented as a schematic illustration in Fig. 10A. Physical observations along with microCT images of nude mice at the point of subcutaneous inoculation showed successful xenograft tumour formation in the WT-OVCAR3 group, with a complete absence of tumours in the shOVCAR3 group (Fig. 10B, C). Excised injection sites after sacrifice corroborated this observation (Fig. 10D, E).

**Fig. 10.**
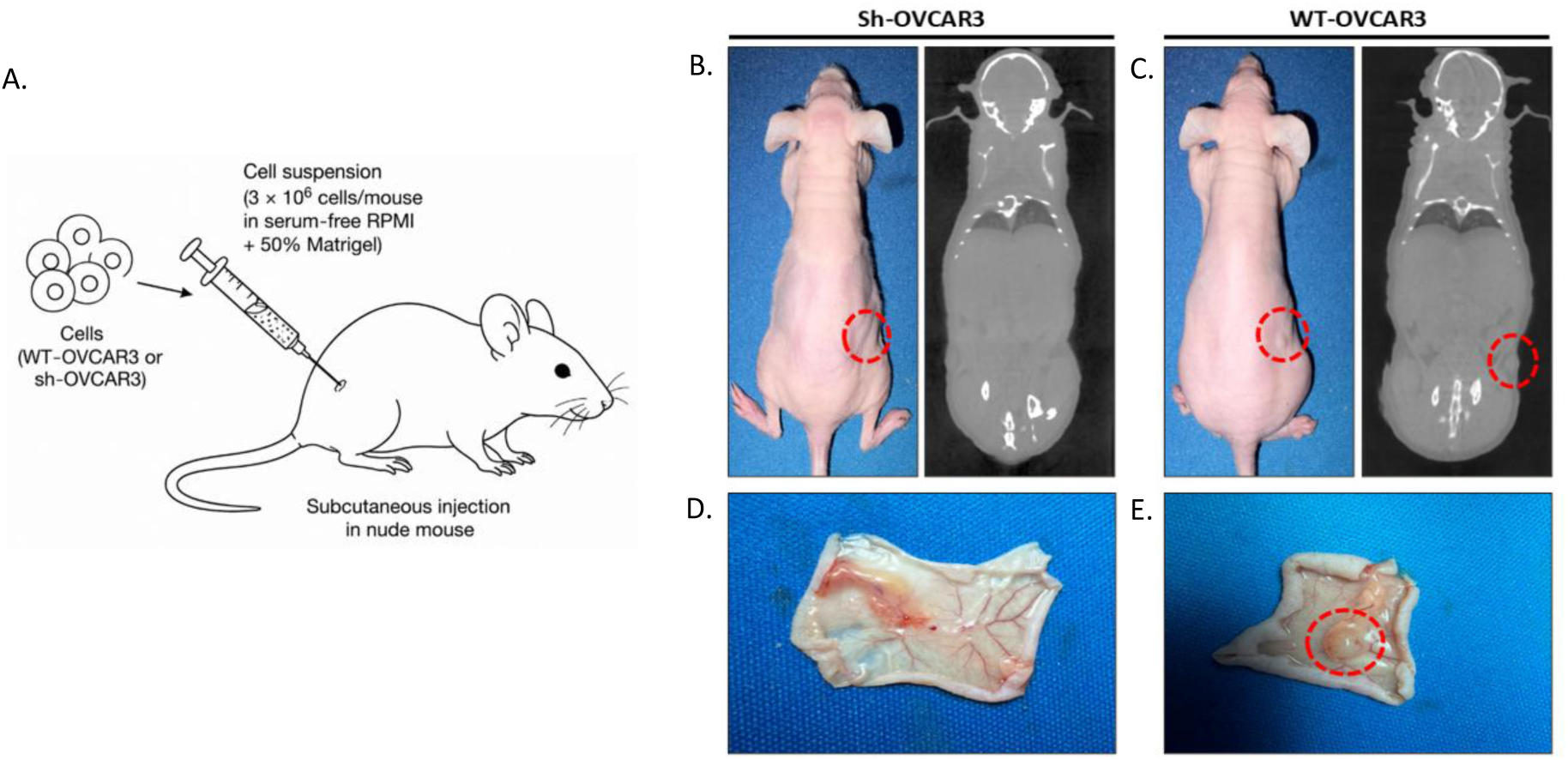
Comparison of tumorigenic potential between sh-OVCAR3 and WT-OVCAR3 cells in a nude mouse xenograft model. **(A)** Schematic illustration of subcutaneous inoculation of WT-OVCAR3 or shOVCAR3 cells into 6–7-week-old nude mice for xenograft tumour establishment. **(B,C)** Representative photographs (left) and corresponding microCT images (right) of nude mice inoculated subcutaneously with shOVCAR3 (B) or WT-OVCAR3 (C) cells, demonstrating the absence of tumour formation in the shOVCAR3 group and successful xenograft tumour formation in the WT-OVCAR3 group (red dashed circles indicate the injection sites). **(D,E)** Gross images of the excised injection sites after sacrifice showing no detectable tumour in the shOVCAR3 group (D), whereas a distinct xenograft tumour in the WT-OVCAR3 group (E, red dashed circles).

Histopathological examination of the injection site from the sh-OVCAR3 group revealed normal dermal and subcutaneous tissue with no evidence of any neoplastic cell proliferation, confirming the absence of tumour formation (Fig. 11, left panel). In contrast, tissue sections from the WT-OVCAR3 group demonstrated a small cellular nodule extending into the dermis and hypodermis (Fig. 10C, E). The tumour was composed of medium- to large-sized round, oval, and polygonal cells arranged in cribriform and solid growth patterns. The neoplastic cells exhibited marked cytonuclear atypia with a high nuclear-to-cytoplasmic ratio. Nuclear pleomorphism was prominent, with nuclei showing round, oval, polygonal, crescent, and stellate shapes. Frequent bizarre nuclei, karyomegaly, and prominent nucleoli were observed, while mitotic figures were infrequent. The nuclear chromatin displayed variable appearances, ranging from sparse granular and hazy to densely compact. Many tumour cells exhibited vesicular cytoplasm. The central region of the tumour showed interstitial oedema, with occasional foci of necrotic debris, whereas minimal peripheral lymphocytic infiltration was observed (Fig. 11, right panel). These histopathological findings reinforced the malignant phenotype and tumourigenic potential of WT-OVCAR3 cells, showing successful engraftment of tumour. This indicates that depletion of lamin A/C expression in sh-OVCAR3 not only affects cellular proliferation and spheroid formation capacity but also induces a lack of tumorigenicity, causing the failure to establish tumours in vivo.

**Fig. 11.**
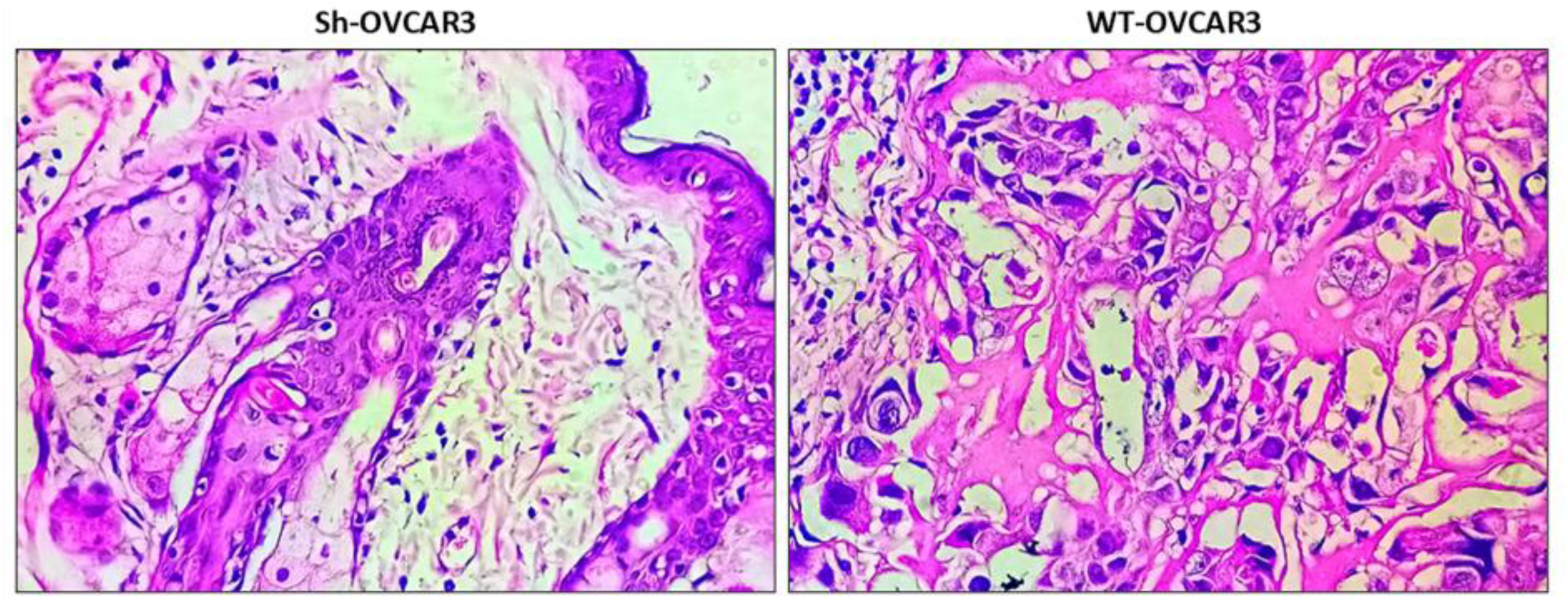
Histopathological evaluation of xenograft tissues derived from sh-OVCAR3 and WT-OVCAR3 cells. Representative haematoxylin and eosin (H&E)-stained sections of the injection sites from shOVCAR3 and WT-OVCAR3 xenografts. The shOVCAR3 group (left) showed normal dermal and subcutaneous tissue architecture without evidence of tumour formation. In contrast, the WT-OVCAR3 group (right) exhibited an infiltrative malignant tumour composed of medium to large-sized pleomorphic cells arranged in cribriform and solid patterns, confirming successful xenograft establishment.

## Discussion

Inside the mammalian nucleus, A-type lamins are ubiquitous in the “nucleographical” parlance (132), and this justifies its interaction with both heterochromatin and euchromatin in health and disease. This interaction specifies a class of euchromatic lamin-associated domains, referred to as A-LADs, that differ from the heterochromatic LADs localised at the nuclear periphery. While classical peripheral LADs involve megabase-scale heterochromatin and are tethered at the nuclear lamina, nucleoplasmic A-LADs scale at finer genomic length and are enriched within more permissive chromatin contexts (9). By dint of stable lamina–chromatin interactions, lamin A/C reduces heterochromatin mobility as well as inhibits transcription largely at the periphery. Genome-wide screening revealed that approximately one-third of lamin A/C-bound loci are localised at the nuclear periphery, whereas the majority transiently reside inside the nucleus (133).

Ovarian carcinoma is primarily associated with increased expression of A-type lamins (36, 107), the consequence of which was previously attributed to increased chemoresistance in HGSOC (39). Like any progressive area of investigation, we scaled vertical depths in exploring the effects of depletion of lamin A/C on tumorigenesis. Not surprisingly, as lamin A/C have long been implicated in modulating 3D chromatin organisation, we followed a similar trail and observed repositioning of chr 18 and 19, representing heterochromatic and euchromatic chromosomes. Similar and rather obvious observations were reported in the case of two heterochromatic chromosomes- 2 and 18 following lamin B1 knockout in breast cancer cell line MDA-MB-231 (134). Our 3D-FISH data found support in the coarse-grained polymer simulation model, which predicted a movement of heterochromatic 18 toward the centre of the nucleus and a complete reversal of trend in the case of euchromatic 19, although the repositioning of chr 19 was less prominent compared to 18. However, we acknowledge some limitations of the present polymer model, which represents single trajectories per condition, generated from a fixed random seed governing the initial chromatin configuration, lamin placement, and stochastic thermal forces. While the wild-type and knockdown trajectories showed a consistent qualitative separation in heterochromatin clustering behaviour, a single realisation cannot distinguish a robust compositional effect from run-to-run stochastic variability inherent to the self-avoiding-walk initialisation and Langevin dynamics.

Nonetheless, this served our purpose. In a sequential flow of events, we asked the next big question: to what extent does this chromosome repositioning following lamin A/C depletion modulate local and long-range chromatin interactions, in particular the local self-interacting topologically associated domains? The differential Hi-C interaction heat map points to a prominent positive interaction profile for chromosome 18, with an increased tendency for active (A) to inactive (B) switching, whereas chromosome19 exhibited higher compartment plasticity with substantial bidirectional toggling and gain of interaction across the off-diagonal matrix consistent with intercompartment mixing. Additionally, we also observed an overall increase in the TAD count as a sequel to this knockdown. In retrospect, integrative analysis of Hi-C in tandem with ChIP-seq was established to dissect super-enhancer elements based on chromatin interactions (135). Super-enhancers are large clusters of adjacent enhancers having unusually high occupancy of interacting factors and have been shown to regulate oncogene expression in cancer (136, 137). The elevated transcriptional output of the cancer cell is brought about by sustained activities of super-enhancer elements (138). Furthermore, super-enhancers and their associated transcription factors are reported to impart chemoresistance in ovarian cancer (139, 140). This prompted us to check the status of such previously reported super-enhancer elements like SE 14, 60, 36, 56 & 86 (67) in our context. For this, we mapped these data with our Hi-C input and found that lamin A/C depletion altered SE-promoter contacts. We supplemented our genomic analyses with RNA sequencing analysis to uncover the transcriptional events that were unfolding. Broadly speaking, we recorded an overall surge in transcriptional activity with major implications in tumorigenesis. The transcriptome could be summarised to predict a significant downregulation of the erythroblastic leukaemia viral oncogene B (ERBB) pathway, which is responsible for cellular proliferation in the context of malignancy. The ERBB represents a family of RTK proteins and comprises of 4 distinct players: the EGFR (ERBB1/HER1), ERBB2 (neu, HER2), ERBB3 (HER3) and ERBB4 (HER4) which can be expressed to different extent and/or in mutated forms in different types of cancer and are involved in a plethora of cellular processes like proliferation, survival, angiogenesis and metastasis thereby rendering them important therapeutic targets (141). Recent advances in ovarian cancer research have shown that the advent of tumour spreading from the ovary and fallopian tube to the peritoneum and abdominal cavities is marked by spheroid formation. This model of peritoneal spread is singularly contributed by ERBB2/EGFR and FOXM1 signalling (131). The diminished expression of EGFR exacerbated the cellular responsiveness towards ligands like TGFA, FGF5, and FGF9, which were all found to be significantly upregulated. The downstream effect percolates into the MAPK and PI3K/AKT signalling axes, which were manifested in an upregulation of SPRY4 and downregulation of DUSP5 and AKT3. It is noteworthy to mention that upregulation of SPRY4 and ASPG bear testimony to restructuring/remodelling of SE 56 and SE 36 and is reminiscent of the B to A transition leading to a surge of transcriptional activity.

In retrospect, highly invasive breast cancer is characterised by an elevation of AKT signalling, which in turn increases nuclear deformation and migratory ability by downregulating lamin A/C (142). We encounter a converse scenario when lamin A/C is depleted, the reason for which is not clearly understood at the moment. Furthermore, due to PTEN depletion in prostate and other cancers, lamin A/C may interact differently with the PI3K/AKT pathway (143–145). It may also be noted that TGF-β-induced EMT leading to chromosomal instability and nuclear deformation is being modulated by AKT2-driven lamin A phosphorylation (146, 147). Therefore, based on previous literature and our findings, we conjecture that lamin A/C perturbs the PI3K/AKT pathway via a two-way traffic which involves complex circuitry and needs further investigation to unravel. We posit that biochemical signalling axes including EGFR, PI3K/AKT and MAPK converge with lamin A/C expression to orchestrate chromatin remodelling, tumour cell invasion, proliferation and survival.

The cumulative observations pertaining to attenuated proliferation in the lamin A/C knockdown scenario were further supported by a significant growth delay in shOVCAR3 compared to WT-OVCAR3. On the other hand, the pronounced reduction in wound closure ability of shOVCAR3 cells is the ex vivo validation of the downregulation of cell motility-associated genes. Finally, xenograft tumour formation from injections of shOVCAR3 in nude mice was largely abrogated, which can also be attributed to the reduction of proliferative potential due to depletion of lamin A/C. Detailed parameters extracted from pathological examinations clearly depicted the fact that shOVCAR3 cells were non-proliferative and non-tumorigenic in nature.

Thus, our research findings have uncovered significant changes in the chromatin landscape when lamin A/C was silenced, which further snowballed into remodelling of super enhancer-promoter interactions and consequent transcriptional output. Differential expression of HGSOC biomarkers like SOX17 and PREX2 as a sequel to lamin A/C depletion opens up newer vistas in exploring the roles of lamin proteins in tumorigenesis. This is but a modest beginning in delineating the roles of lamin A/C in the potentiation of HGSOC. We acknowledge that miles of uncharted territory are therefore left to be explored in terms of the interaction of lamin A/C with some of these important proteins. The physiological levels of lamin A/C aid in organising the 3D genome, conferring chromatin flexibility and stability, resulting in cellular homeostasis. In cancer, these are perturbed in favour of abnormal proliferation, migration and other biochemical pathways that go awry. LMNA, though not being an oncogene or tumour suppressor, modulates genomic stability and subsequent downstream processes like a nuclear “master regulator”. Lamin A/C, being a transcription modulator, has immense possibilities of directly or indirectly driving the promoters of those genes implicated in the proliferation and progression of HGSOC or simply can induce the bystander effect by complex mechanisms hitherto unknown. This report thus outlines the global effects of lamin A/C depletion on chromatin reorganisation and subsequent hyperfine topological elements, thereby affecting gene regulation. We demonstrated that lamin A/C depletion can affect both euchromatic and heterochromatic chromosomes, unlike lamin B depletion, which translates primarily into heterochromatin disorganisation (134). Future research in the area of lamin A/C mediated pathogenesis of HGSOC is likely to shed more light on the diverse roles of lamin A/C as the guardian of the nucleus. Therefore, it would not be an overstatement to predict tunable interventions of lamin A/C levels to control the signalling axes leading to neoplastic transformations.

## Supporting information

OV_Sh.cis.vecs

OV_sh_50kb_raw.cool

OV_WT_50kb_raw.cool

OV_WT.cis.vecs

OVCAR3_superenhancers.bed

SE14_SE60_promoter_contact_comparison_50kb

OVCAR3_SE_compartment_transitions

## Acknowledgements

We acknowledge Ms Trishita Banerjee, ACTREC, Tata Memorial Centre, Mumbai, India, for helping with the preparations for animal experiments in the initial phase.

## Author contributions

SDS performed all wet lab experiments and some bioinformatic analysis and wrote the manuscript; PG performed bioinformatic analysis and wrote the respective parts of the manuscript; AP, SP performed coarse grained simulations wrote the respective parts of the manuscript; VSM, BSM performed animal experiments and wrote the respective parts of the manuscript; RB performed the Hi-C experiments; AB supervised the coarse grained modelling studies; PC supervised the animal experiments; KB facilitated animal experiments; KSG designed and conceptualized the whole project, collated all data and wrote the manuscript.

## Supplementary Data

1. **Supplementary file 1**
2. **Supplementary file 2**
3. **OVCAR3_SE_compartment_transitions.tsv**
4. **SE14_SE60_promoter_contact_comparison_50kb.tsv**
5. **OVCAR3_superenhancers.bed**
6. **OV_WT_50kb_raw.cool**
7. **OVCAR3_SE_compartment_scores.tsv**
8. **OV_WT.cis.vecs.tsv**
9. **OV_Sh_50kb_raw.cool**
10. **OV_Sh.cis.vecs.tsv**
11. **Supplementary_Materials_&_Methods_and_figures_Final_version**

## Conflict Of Interest

Authors contributing to the manuscript declare no conflict of interest.

## Funding

This project is funded from the intramural project RSI 4002, Department of Atomic Energy, Govt. of India

## Data Availability

The Hi-C and RNA-sequencing data generated in this study have been deposited in the NCBI Gene Expression Omnibus (GEO) under accession numbers GSE345872 (Hi-C): https://www.ncbi.nlm.nih.gov/geo/query/acc.cgi?acc=GSE345872 and GSE345919 (RNA-seq): https://www.ncbi.nlm.nih.gov/geo/query/acc.cgi?acc=GSE345919.

## Code Availability

1. The simulation code used in this study is publicly available at: https://github.com/cosmicpanda1/MD-simulation
2. All other Hi-C, and RNA sequencing datasets supporting the findings of this study and the codes used are available within the article and its Supplementary Information or from the corresponding author upon reasonable request.

## Notes

### Competing Interest Statement

The authors have declared no competing interest.

