## Supplementary_Materials_&_Methods_and_figures for "Remodelling of chromatin architecture and super-enhancer landscape in lamin A/C depleted HGSOC"

### RNA-Seq pathway analysis

Functional enrichment analysis was performed using the clusterProfiler package. Significantly upregulated and downregulated genes were analysed separately using Gene Ontology (GO) Biological Process pathway enrichment. Enrichment analysis was performed using Entrez Gene IDs. Statistical significance was assessed using the hypergeometric test, followed by Benjamini–Hochberg correction for multiple testing. GO terms with an adjusted p-value (FDR) $< 0.05$ were considered significantly enriched. To facilitate biological interpretation, enriched GO terms were further annotated by identifying the differentially expressed genes contributing to each enriched biological process together with their corresponding $\log_2$fold-change values. To investigate coordinated changes in biological pathways without applying an arbitrary fold-change threshold, Gene Set Enrichment Analysis (GSEA) was performed using the fgsea package. Genes were ranked according to the DESeq2 Wald statistic, preserving both the magnitude and direction of differential expression between shOVCAR3 and WT-OVCAR3 samples. Positive ranking scores corresponded to genes more highly expressed in shOVCAR3, whereas negative ranking scores corresponded to genes more highly expressed in WT. Gene sets were obtained from the Molecular Signatures Database (MSigDB) (1) using the msigdbr package. Three collections were analysed: Hallmark gene sets (H collection), Reactome pathways (C2: CP: REACTOME), and Gene Ontology Biological Process gene sets (C5: GO: BP). Enrichment scores were calculated using the fgsea algorithm, and statistical significance was determined after false discovery rate correction. Pathways with an adjusted p-value $< 0.05$ were considered significantly enriched. Positive normalised enrichment scores (NES) indicate enrichment toward genes more highly expressed in shOVCAR3 samples, whereas negative NES values indicate enrichment toward genes more highly expressed in WT-OVCAR3 samples. Leading-edge analysis was used to identify the subset of genes contributing most strongly to each enriched pathway. All steps were performed in R version 4.5.3.

To investigate the ERBB-associated Reactome pathway analysis, the Reactome GSEA results were filtered for terms containing ERBB to extract ERBB-related pathways. Ten such pathways were identified, and the GSEA leading genes were extracted for individual pathways. The resulting pathway-gene table contained 94 genes, out of which only 27 unique genes were obtained for subsequent network analysis. Pathway association was retained independently so that genes appearing in more than one ERBB-related pathway could be distinguished. The pathways with FDR-adjusted p-value $< 0.05$ were considered significantly enriched.

For STRING interaction-network analysis, the 27 unique ERBB-related genes were queried against the STRING interaction database to identify functional associations. A high-confidence STRING interaction threshold of 0.70 was applied, and interactions among the input gene set were considered. The final interaction network was visualised using NetworkX and Matplotlib.

**Chromosome-specific A/B compartment analysis and integration with transcriptomic data**

A/B compartments were identified from balanced 1 Mb Hi-C contact matrices using the eigs-cis function implemented in cooltools, which performs principal component analysis (PCA) on observed/expected contact matrices to generate the first eigenvector (E1). Since the orientation of PCA eigenvectors is mathematically arbitrary, the direction of the E1 values was validated biologically using genome-wide genomic features, including GC content and gene density. Genome-wide GC content for each 1 Mb genomic bin was obtained using the hg38 reference genome and compared with the corresponding E1 values. Gene density was independently assessed by assigning all annotated genes from the GENCODE v49 primary assembly annotation to their respective 1 Mb genomic bins based on the transcription start site (TSS), followed by calculation of the number of genes per bin. Following these validations, the eigenvector orientation was uniformly adjusted so that positive E1 values represented transcriptionally active A compartments and negative E1 values represented inactive B compartments. The final oriented E1 values were extracted for chromosomes 18 and 19 for visualisation of chromosome-level compartment organisation. Identical genomic coordinates and a common y-axis were used to facilitate direct comparison between conditions. For genome-wide compartment analysis, each 1 Mb genomic bin was classified according to its compartment status in WT and Lamin-knockdown cells as A→A, B→B, A→B or B→A. Changes in compartment organisation were quantified using the difference in compartment eigenvector values ($\Delta E1 = E1_{shOVCAR3} - E1_{WT\text{-}OVCAR3}$).

To check whether compartment remodelling is associated with transcriptional changes, RNA-sequencing differential expression results were integrated with compartment assignments. Genes were mapped to their corresponding 1 Mb compartment bins according to their TSS coordinates. Differentially expressed genes were defined using an adjusted p-value $< 0.05$ together with an absolute $\log_2$ fold change $\geq 0.585$ (equivalent to a minimum 1.5-fold expression change). Each gene was subsequently assigned to the compartment transition of its genomic bin, allowing comparison of transcriptional changes among stable and switching compartment classes. Differences in expression distributions were evaluated using Mann–Whitney U tests, and genome-wide associations between compartment remodelling ($\Delta E1$) and transcriptional change were assessed using Spearman rank correlation.

**Super-enhancer loci analysis from Hi-C data**

86 candidate super-enhancer (SE) loci were selected to be studied from an earlier reference (2). The genomic coordinates of the reported OVCAR3 SE loci were extracted from the supplementary dataset: https://www.nature.com/articles/s41467-022-31919-8#MOESM3 and used as a reference to map onto Hi-C datasets generated from WT-OVCAR3 and shOVCAR3 cells in the present study to assess their chromatin organisation following Lamin depletion. The

SEs were targeted by two distinct sgRNA designs (A and B), where the full SE genomic interval for each locus was reported on the A-designated row. Therefore, A-designated rows were extracted to represent one non-redundant genomic coordinate per SE and were converted from 1-based genomic notation to 0-based BED format by subtracting one from the start coordinate while retaining the exact end coordinate (Supplementary file, OVCAR3_superenhancers.bed). The published SE coordinates were intersected with A/B compartment profiles with the Hi-C data generated from WT-OVCAR3 and shOVCAR3 cells. Cooltools eigs-cis was used to obtain compartment profiles from 1-Mb Cooler contact matrices. Eigenvector orientation was phased using a 1-Mb hg38 GC-content track so that the positive E1 values corresponded to the transcriptionally active and GC-rich compartment A and the negative E1 values corresponded to the B compartment (Supplementary files, OV_WT.cis.vecs.tsv and OV_Sh.cis.vecs.tsv). Each SE belongs to a 1-Mb compartment bin containing the respective genomic midpoint, and the SE locus is classified as stable A (A→A), stable B (B→B), B-to-A switching (B→A) or A-to-B switching (A→B) for both cases, WT-OVCAR3 and shOVCAR3. Loci that cannot be assigned to a valid compartment were classified as unassigned. The E1 values associated with each SE were also compared between the states to quantify changes in compartment strength.
The respective coordinates SE14, chr1:16,493,670-16,516,686 and SE60, chr20:52,352,050-52,368,920. Target gene transcription start sites were obtained from the GENCODE v49 primary-assembly basic annotation gtf file. The promoter-containing Hi-C bins were identified from the transcription start sites according to gene strand. SE-promoter chromatin interactions were assessed with WT-OVCAR3 and shOVCAR3 50-kb Cooler matrices (Supplementary files: OV_WT_50kb_raw.cool and OV_Sh_50kb_raw.cool). Matrix bins and stored balancing weights were accessed to extract balanced intrachromosomal Hi-C contact matrices using Cooler (3). To include the distance-dependent decay of Hi-C contact frequency, observed/expected (O/E) interaction matrices were calculated for each chromosome and condition. For each genomic bin separation, the mean of all valid (finite and non-zero) balanced contacts along the corresponding matrix (chromosome) diagonal was used as the expected contact frequency. Each observed balanced contact was then divided by the distance-matched expected value for the same genomic separation to obtain the O/E interaction frequency.
For each SE-promoter pair, the bin containing the target-gene transcription start site and the 50-kb bin containing the SE midpoint were identified. Interaction strength was calculated as the mean O/E contact frequency within a 3 x 3 bin window centred on the promoter and SE anchor bins, including the anchor interaction and the adjacent 50-kb bins. WT-OVCAR3 and shOVCAR3 values were compared using the O/E ratio (shOVCAR3/WT-OVCAR3) and percentage change. Regional $\log_2$(O/E) contact matrices were visualised using Python scripts with identical colour scales between conditions, and the SE and promoter anchor positions were annotated on the Hi-C maps. The resulting SE-level compartment and chromatin-interaction analyses were generated using Python scripts (Supplementary files, OVCAR3_SE_compartment_transitions.tsv, OVCAR3_SE_compartment_scores.tsv and SE14_SE60_promoter_contact_comparison_50kb.tsv). Using differential expression results, the DESeq2-derived $\log_2$ fold changes and Benjamini–Hochberg-adjusted P values for EPHA2 and RAE1 were extracted using gene symbols. These transcriptional measurements were then integrated with the corresponding SE-promoter O/E contact changes to assess whether changes in SE–promoter contact strength were accompanied by changes in target-gene expression in shOVCAR3 with respect to WT-OVCAR3.

**FENE bonding potential**

To maintain polymer integrity and prevent bond breaking, adjacent chromatin beads were connected using a finitely extensible nonlinear elastic (FENE) potential (4):

$$U_{FENE}(r) = \frac{-1}{2} k_{FENE} R_0^2 ln\left[1 - \left(\frac{r}{R_0}\right)^2\right],$$

where $k_{FENE} = 30k_B T/\sigma^2$ is the spring constant and $R_0 = 1.6\sigma$ is the maximum bond extension. The corresponding FENE bond force is

$$F_{FENE}(r) = \frac{k_{FENE} r}{1 - (r/R_0)^2} \hat{r},$$

directed along the bond vector between adjacent chromatin beads, and similarly capped at $F_{max}$ to prevent numerical divergence as $r \to R_0$. This potential ensured strong connectivity at short distances while preventing non-physical bond stretching beyond $R_0$. Compared to the harmonic springs, the FENE formulation, which is widely used in coarse-grained polymer simulations (4), provided enhanced numerical stability.

**Interaction matrix and bonding potentials**

Different affinities between lamin A (LA), lamin B (LB), heterochromatin (HC), and euchromatin (EC) were allowed by parameterising all non-bonded contacts using an explicit interaction matrix derived from the simulation code (Table 1). For every pair of species, the interaction strengths $\epsilon_{ij}$ and matching cutoff distances $r_{c,ij}$ were computed. While heterochromatin demonstrated substantial self-attraction ($\epsilon_{HC,HC} = 2.0$), encouraging dense phase growth, euchromatin showed modest self-attraction ($\epsilon_{EC,EC} = 0.2$), indicating its open and less compact structure. The intermediate cross-interactions between heterochromatin and euchromatin ($\epsilon_{EC,HC} = 1.0$) allowed for partial mixing (5, 6). Lamin-chromatin interactions were tuned to capture differential binding affinities. Euchromatin interacted moderately with lamin A ($\epsilon_{EC,L} = 0.75$) and weakly with lamin B ($\epsilon_{EC,LB} = 0.30$). In contrast, heterochromatin exhibited stronger affinity towards both lamin species under wild-type conditions, with $\epsilon_{HC,L} = 1.25$ and $\epsilon_{HC,LB} = 2.0$, promoting peripheral localisation and stabilisation of condensed domains (7, 8).

**Table 1.** *Interaction strengths$\varepsilon_{ij}$ (in units of $k_B T$) between euchromatin (EC), heterochromatin (HC), lamin A (L), and lamin B (LB), as used in the wild-type (WT) simulations. Values for heterochromatin–lamin interactions ($\varepsilon_{HC,L}$, $\varepsilon_{HC,LB}$) and lamin self-interactions ($\varepsilon_{L,L}$, $\varepsilon_{LB,LB}$) are condition-dependent; WT values are shown here (see Methods for the corresponding knockdown parameterisation).*

| | EC | HC | L | LB |
|---|---|---|---|---|
| EC | 0.2 | 1.0 | 0.75 | 0.30 |
| HC | 1.0 | 2.0 | 1.25 | 2.0 |
| L | 0.75 | 1.25 | 1.0 | 0.33 |
| LB | 0.30 | 2.0 | 0.33 | 0.5 |

The cutoff distances were scaled according to particle sizes with the following criterion (Table 2):

$$r_{c,ij} \quad \alpha_{ij}\frac{\sigma_i + \sigma_j}{2},$$

where $\alpha_{ij}$ is an interaction-specific scaling factor and $\sigma_{ij} = (\sigma_i + \sigma_j)/2$ is the mean bead diameter of the pair.

The character of each pairwise interaction was found to be purely repulsive (steric) or attractive, as set by the ratio of the cutoff distance $r_{c,ij}$ to the Weeks-Chandler-Andersen (WCA) threshold $2^{1/6}\sigma_{ij}$. When $r_{c,ij} \leq 2^{1/6}\sigma_{ij}$, the potential was truncated before reaching its attractive minimum so that the interaction was purely repulsive (excluded volume only) regardless of the magnitude of $\epsilon_{ij}$; when $r_{c,ij} > 2^{1/6}\sigma_{ij}$, the interaction retained an attractive well.

**Table 2.** *Cutoff scaling factors $\alpha_{ij}$, resulting cutoff distances $r_{c,ij}$, and interaction character for each particle pair. The WCA threshold is $2^{1/6}\sigma_{ij}$, where $\sigma_{ij} = (\sigma_i + \sigma_j)/2$; pairs with $r_{c,ij}$ below this threshold are purely repulsive (steric), while pairs above it retain an attractive well.*

| Pair | $\alpha_{ij}$ | $\sigma_{ij}$ | $r_{c,ij}$ | Character |
|---|---|---|---|---|
| EC–EC | 1.80 | $1.0\sigma$ | $1.80\sigma$ | Attractive |
| HC–HC | 1.80 | $1.0\sigma$ | $1.80\sigma$ | Attractive |
| EC–HC | 1.12 | $1.0\sigma$ | $1.12\sigma$ | Repulsive (WCA) |
| EC–L | 0.84 | $0.75\sigma$ | $0.63\sigma$ | Repulsive (WCA) |
| EC–LB | 0.80 | $0.70\sigma$ | $0.56\sigma$ | Repulsive (WCA) |
| HC–L | 1.35 | $0.75\sigma$ | $1.0125\sigma$ | Attractive |
| HC–LB | 1.35 | $0.70\sigma$ | $0.945\sigma$ | Attractive |
| L–L | 0.60 | $0.50\sigma$ | $0.30\sigma$ | Repulsive (WCA) |
| LB–LB | 0.60 | $0.40\sigma$ | $0.24\sigma$ | Repulsive (WCA) |
| L–LB | 0.50 | $0.45\sigma$ | $0.225\sigma$ | Repulsive (WCA) |

Self-interactions among euchromatin ($\alpha = 1.8$) and heterochromatin ($\alpha = 1.8$) both retained a full attractive well, consistent with the compaction propensities described above. In contrast, euchromatin-heterochromatin cross-interactions used a WCA-truncated cutoff ($\alpha = 1.12$, i.e. $r_{c,EC,HC} \approx 2^{1/6}\sigma$) and were hence purely repulsive. It is to be noted that in conjunction with the strong self-attraction of heterochromatin, this repulsive cross-term is the primary driver of separation between the two chromatin states, rather than any attractive coupling between them. Lamin-euchromatin cross-interactions ($\alpha = 0.84$ for EC-L, $\alpha = 0.80$ for EC-LB) were likewise WCA-truncated and purely repulsive, acting only to exclude volume between euchromatin and lamin particles. Heterochromatin-lamin interactions, by contrast, used longer cutoffs ($\alpha = 1.35$ for both HC-LA and HC-LB), retaining an attractive well and providing the effective driving force for heterochromatin recruitment toward lamin-associated regions. Lamin self-interactions (L-L and LB-LB, $\alpha = 0.6$) and the lamin A-lamin B cross-interaction (LA-LB, $\alpha = 0.5$) were all WCA-truncated and purely repulsive. Direct pairwise lamin-lamin attraction therefore did not contribute to lamin clustering in this model. As described in the following subsection, lamin accumulation at the nuclear periphery instead arose from an explicit confining wall potential.

**Confining boundary and nuclear envelope potential**

Periodic boundary conditions were applied in the lateral (x, y) directions only. The axial (z) direction was bounded by confining walls at $z_{\min} = -L_z/2$ and $z_{\max} = +L_z/2$, representing the nuclear envelope.

Chromatin beads (euchromatin and heterochromatin) experienced a weak, purely repulsive WCA-type wall potential at both boundaries, preventing escape from the simulation domain without imposing any preferential localisation.

Lamin A and lamin B particles experienced an asymmetric wall potential: an attractive interaction with the upper wall ($z_{\max}$), of strength $\epsilon_{ML}$=8.5 for lamin A and $\epsilon_{MLB}$=15 for lamin B, and a substantially weaker attraction $(0.1 \times \epsilon_{\text{ML}}, 0.1 \times \epsilon_{\text{MLB}})$ with the lower wall at $z_{\min}$. Each wall interaction was implemented as a truncated Lennard–Jones-type potential acting on the distance z between a particle and the relevant wall,

$$U_{\text{wall}}(z) = 4\epsilon_w \left[ \left( \frac{\sigma_t}{z} \right)^{12} - \left( \frac{\sigma_t}{z} \right)^{6} \right],$$

where $\sigma_t$ is the diameter of the bead of the particle type $t$ and $\epsilon_w$ is the interaction strength between the type and the wall. For chromatin beads (EC, HC), $\epsilon_w = 0.1\,\epsilon_{\text{EC,EC}}$ on both walls, giving a weak, purely repulsive confinement. For lamin A and lamin B, $\epsilon_w = \epsilon_{\text{ML}}$ or $\epsilon_{\text{MLB}}$ on the upper wall ($z_{\max}$) and 0.1 $\epsilon_{\text{ML}}$ or 0.1 $\epsilon_{\text{MLB}}$ on the lower wall ($z_{\min}$), respectively, producing the asymmetric peripheral attraction described above. This asymmetric wall attraction represents the preferential association of lamin proteins with the nuclear envelope and is the principal mechanism driving lamin peripheral localisation in the model, given that direct lamin-lamin pairwise interactions are purely repulsive, as shown above. This wall-based representation of the nuclear lamina contrasts with discrete-particle lamin models (8) and earlier surface-layer formulations (7, 9).

Supplementary Figures:

Sup Fig.1

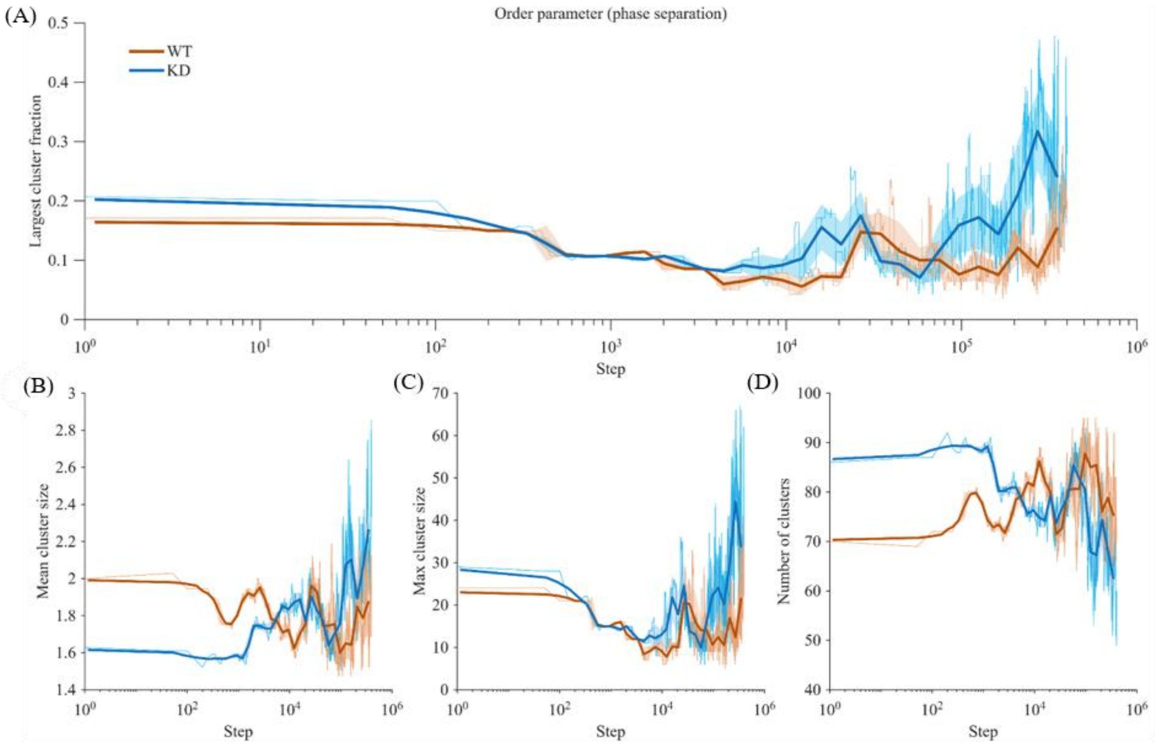


Sup. Fig.1. **Log-binned time evolution of heterochromatin (HC) clustering in WT-OVCAR3 and shOVCAR3 simulations.** Simulation-derived quantification of heterochromatin clustering dynamics over a 4x10^5-step trajectory comparing wild-type (WT, orange) and lamin A/C knockdown (KD, blue) conditions: (A) Order parameter dynamics, defined as the fraction of total HC beads contained within the largest cluster. (B) Mean HC cluster size, (C) Largest HC domain (maximum cluster size attained), (D) total number of HC clusters. Simulation trajectory length is expressed in units of time steps. Thin curves represent raw, per-step values throughout the trajectory, while bold curves and shaded bands denote the mean and standard deviation within logarithmically spaced step bins, capturing temporal variability within-trajectory.

Sup Fig.2

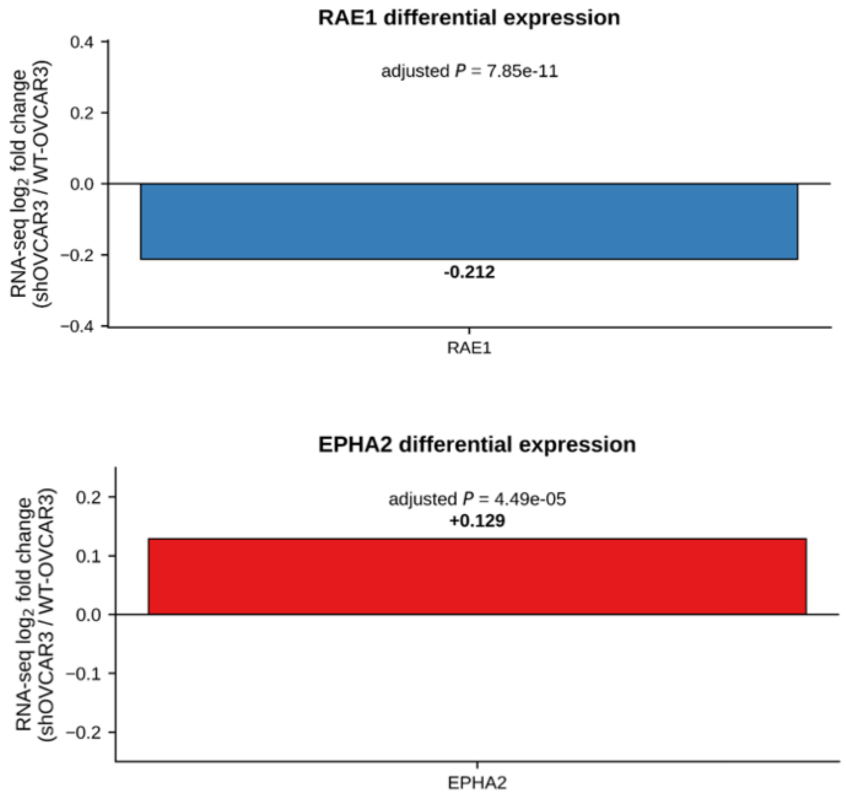


Sup. Fig.2. **Quantification of differential expression of transcripts in shOVCAR3 relative to WT-OVCAR3.** (A) RNA-seq differential-expression result for RAE1, shown as the $\log_2$ fold change. (B) RNA-seq differential-expression result for EPHA2 in terms of $\log_2$ fold change
